# Cytoplasmic polyadenylation is an ancestral hallmark of early development in animals

**DOI:** 10.1101/2023.05.10.540224

**Authors:** Labib Rouhana, Allison Edgar, Fredrik Hugosson, Valeria Dountcheva, Mark Q. Martindale, Joseph F. Ryan

## Abstract

Differential regulation of gene expression has produced the astonishing diversity of life on Earth. Understanding the origin and evolution of mechanistic innovations for control of gene expression is therefore integral to evolutionary and developmental biology. Cytoplasmic polyadenylation is the biochemical extension of polyadenosine at the 3’-end of cytoplasmic mRNAs. This process regulates the translation of specific maternal transcripts and is mediated by the Cytoplasmic Polyadenylation Element Binding Protein family (CPEBs). Genes that code for CPEBs are amongst a very few that are present in animals but missing in non-animal lineages. Whether cytoplasmic polyadenylation is present in non bilaterian animals (*i.e.*, sponges, ctenophores, placozoans, cnidarians) remains unknown. We have conducted phylogenetic analyses of CPEBs and our results show that CPEB1 and CPEB2 subfamilies originated in the animal stem lineage. Our assessment of expression in the sea anemone, *Nematostella vectensis* (Cnidaria), and the comb jelly, *Mnemiopsis leidyi* (Ctenophora), demonstrates that maternal expression of CPEB1 and the catalytic subunit of the cytoplasmic polyadenylation machinery (GLD2) is an ancient feature that is conserved across animals. Furthermore, our measurements of poly(A)-tail elongation reveal that key targets of cytoplasmic polyadenylation are shared between vertebrates, cnidarians, and ctenophores, indicating that this mechanism orchestrates a regulatory network that is conserved throughout animal evolution. We postulate that cytoplasmic polyadenylation through CPEBs was a fundamental innovation that contributed to animal evolution from unicellular life.

## INTRODUCTION

Post-transcriptional control of gene expression encompasses events that influence mRNA localization, stability, and rate of translation [1–3]. While gene expression can be regulated at many levels, post-transcriptional regulation of mRNA is known to be particularly relevant during early stages of animal development [4–9], where the initial zygotic cleavages can occur from cytoplasmic material deposited in the egg and in the absence of nuclear components, including chromosomal DNA [10, 11]. Instead of being driven by transcriptional events, changes in gene expression during ovulation and immediately after fertilization are orchestrated through cytoplasmic regulation of maternal transcripts. Post-transcriptional regulation of maternal mRNA results in timely production of proteins that control cell cycle progression and cell fate determination during early development [7, 12–14].

Cytoplasmic polyadenylation is a post-transcriptional mechanism for regulating gene expression that involves lengthening of the 3’-polyadenosine tail of mRNA (*i.e.,* the poly(A)-tail) following nuclear polyadenylation and export from the nucleus [3, 4, 15–19]. Lengthening the poly(A)-tail through this mechanism correlates with stability and higher translational activity, whereas its shortening leads to dormancy and decay [20–22]. In contrast to canonical polyadenylation events that take place during transcriptional termination in the nucleus, which are part of the usual processing for the vast majority of eukaryotic mRNAs, cytoplasmic polyadenylation is limited to specific substrates and biological contexts [16, 23]. This phenomenon was first documented, to our knowledge, during experimentally-induced cleavage of enucleated sea urchin embryos [24]. In the time since, studies of oogenesis and early embryonic development in a handful of bilaterian animal models have identified homologs of the Cytoplasmic Polyadenylation Element-Binding Protein (CPEB) as conserved regulators for cytoplasmic polyadenylation of maternal mRNAs [16, 17, 25]. The best-characterized cytoplasmic polyadenylation complexes contain three primary components: a CPEB, a non-canonical poly(A) polymerase (*e.g.,* GLD2), and subunits of the Cleavage and Polyadenylation Specificity Factor (CPSF) (Figure 1A). CPEBs provide substrate specify to the complex [26, 27], while members of the GLD-2 family provide enzymatic activity [28–31]. Cytoplasmic polyadenylation not only requires CPSF subunits but also a sequence element that these recognize during transcriptional termination (the hexanucleotide AAUAAA; [32, 33]), which because of its ubiquitous presence in mRNA contributes negligibly to target selection in the cytoplasm.

**Figure 1.**
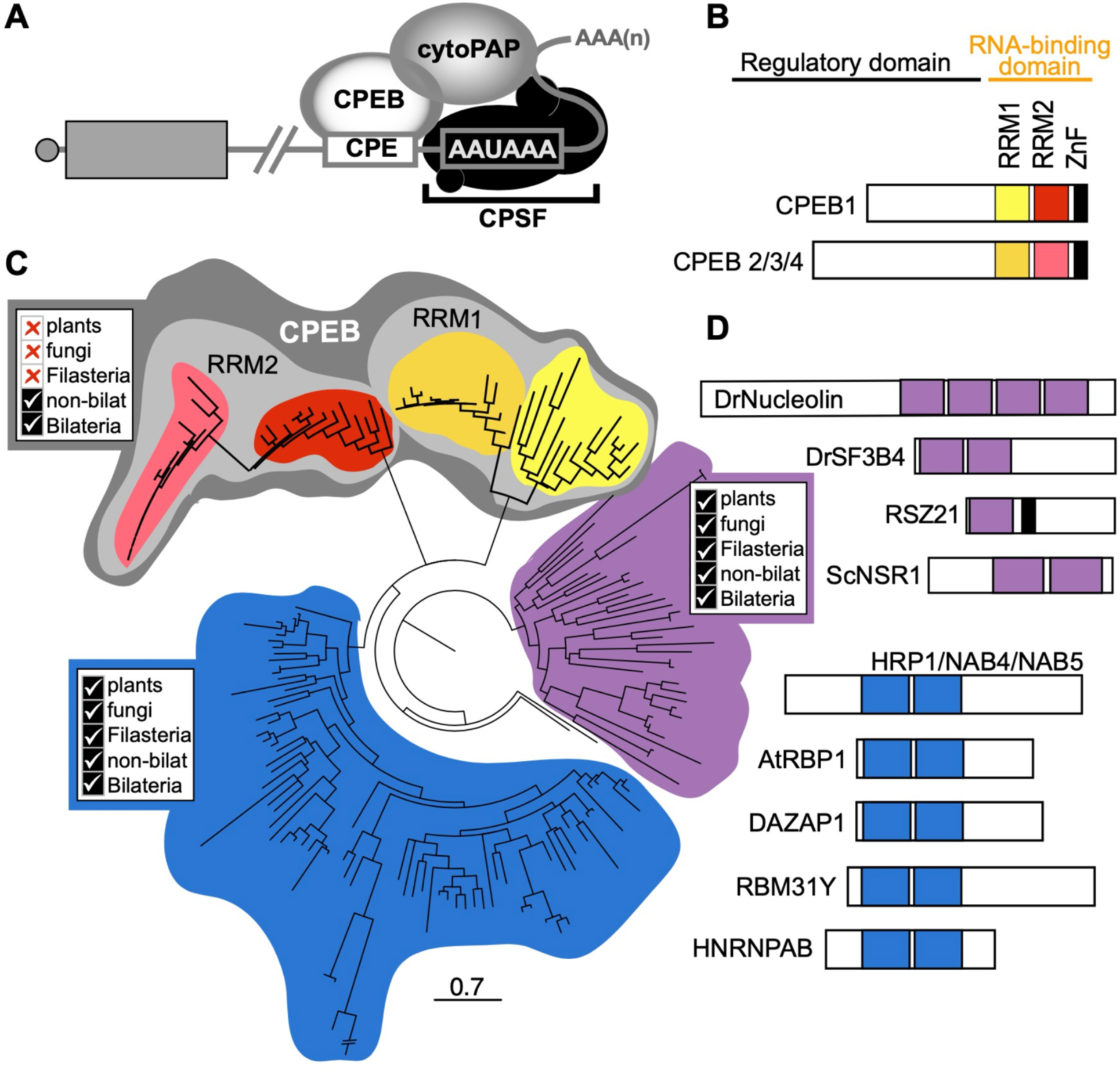
Cytoplasmic Polyadenylation Element Binding Proteins (CPEBs) are highly conserved post transcriptional regulators present at the stem of animal evolution. **(A)** Diagram depicting the core cytoplasmic polyadenylation complex. Subunits of the Cleavage and Polyadenylation Specificity Factor (CPSF) are bound to the hexanucleotide A2UA3, and a CPEB bound to a Cytoplasmic Polyadenylation Elements (CPE) recruits a poly(A) polymerase of the GLD2 family (cytoPAP) to the 3’-end of mRNAs. **(B)** CPEBs contain a highly conserved C-terminal RNA-binding domain composed of two RNA-Recognition Motifs (RRMs; yellow for RRM1 and red for RRM2 of CPEB1; orange for RRM1 and pink for RRM2 of CPEB2/3/4 subfamily members) and a Zinc-Finger motif (ZnF; black). The N-terminus of CPEBs is not well conserved but is known to contain regulatory elements. Diagram based on representatives from *S. mediterranea* [38]. **(C)** Maximum likelihood phylogenetic tree based on similarity to individual RRMs of human CPEB1 depicts relationship between CPEB orthologs and close homologs. The first (RRM1) and second (RRM2) RNA-Recognition Motifs of previously characterized CPEBs (*e.g., Mus musculus* CPEB1-4) positioned in separate clades (light gray shading) that were only composed of metazoan sequences (color coding for sub-clades correspond to those used for respective RRMs in panel (B). Proteins from non-animal species, such as plant, fungi, and choanoflagellates, are only found in clades that lacked CPEBs (shaded in blue and purple). **(D)** Bar diagrams of representative proteins in clades with homology to RRMs of CPEBs. RRMs are depicted in the color shading of their corresponding clades in panel (C) and ZnF shown in black. Scale bar represents substitutions per amino acid position.

CPEBs are divided into two major subfamilies. Members of the CPEB1 subfamily are required for oogenesis and regulation of maternal mRNAs during early development across Bilateria [26, 34–38]. For example, mutations in the *Drosophila* CPEB1 ortholog *oo18 RNA binding* (*orb*) obstructs oogenic progression during (null) and after (*orb^303^*) 16-cell cyst formation, whereas a less-severe mutation (*orb^mel^*) results in abnormal embryonic development due to misregulation of localized mRNA translation [39–41]. Members of the CPEB2 subfamily (*e.g.* CPEB2, CPEB3, and CPEB4 in mice and humans; Orb2 in *Drosophila*; and CPB-1 and FOG-1 in *C. elegans*) display enriched expression in animal testes and some are known to be required for sperm development [38, 42, 43]. Similarly, homologs of the catalytic subunit of the cytoplasmic polyadenylation complex (*i.e.,* the GLD2 family of non-canonical poly(A) polymerases) are also required for oogenesis, spermatogenesis, and early embryonic development [28, 30, 31, 44–46]. Outside of the germline, CPEBs and GLD2 homologs have a conserved role in regulating localized translation of neuronal transcripts, and their function is required for memory formation and synaptic plasticity [47–54]

The C-terminus of CPEBs contains a highly conserved RNA-Binding Domain (RBD), which is composed of two RNA-Recognition Motifs (RRMs) and a Zinc Finger (Figure 1B). The RBD of CPEBs from bilaterian phyla (Deuterostomia, Lophotrochozoa, and Ecdysozoa) contain primary structure that is more than 30% identical (Rouhana et al., 2017), whereas the rest of the protein lacks recognizable sequence conservation (Figure 1B; reviewed by [55]). CPEB1 orthologs bind to well-defined Cytoplasmic Polyadenylation Elements (CPEs; consensus sequence: UUUUA(U/A)) present in the 3’UTR of their targets, which include *c-mos*, *cyclin*, and *Dazl* mRNAs [56–62]. There is less conservation or agreement regarding sequence recognition by members of the CPEB2 subfamily in comparison to that of CPEB1, but these also seem to have affinity for U-rich elements [63–65]. Evidence exists for co-regulation of some targets of cytoplasmic polyadenylation by members of both CPEB subfamilies [60, 66–68], as well as for functions for CPEBs that are independent of modulation of poly(A)-tail length [69–72]. However, the majority of studies show that the central role of CPEBs is in regulating specific subsets of mRNA via cytoplasmic regulation of poly(A) tail length [15, 16, 25, 73–76].

Despite its important regulatory role in animal development, it is not known whether cytoplasmic polyadenylation is present in non-bilaterian animals. Homologs of GLD2 and CPSF subunits are present across Eukarya and involved in nuclear processes [77–79]. However, a recent global survey determined that CPEB homologs are absent outside of animals and present in genomes from every major animal lineage (see Supplementary Table S1) [80]. These findings are consistent with the hypothesis that cytoplasmic polyadenylation arose in the lineage leading to the last common ancestor of animals. However, the results relied on similarity-based methods and require phylogenetic confirmation as well as functional evidence. Here we phylogenetically analyze the presence of CPEB1 and CPEB2 subfamily members in genomes of species that belong to each of the five major animal lineages, which are bilaterians (that account for 99% of all extant animal species) and the four earlier branching non-bilaterian clades (ctenophores, placozoans, sponges, and cnidarians), as well as in genomes of non-metazoan models. We also assess whether CPEBs and other members of the cytoplasmic polyadenylation complex are maternally deposited in eggs of the cnidarian model *Nematostella vectensis* and the ctenophore *Mnemiopsis leidyi.* Finally, we determine whether CPEB targets identified in studies of maternal mRNA regulation in vertebrate models display conserved changes in poly(A)-tail length during oocyte maturation and early embryonic development in cnidarians and ctenophores. Our findings suggest that CPEB-mediated cytoplasmic polyadenylation is an ancestral mechanism that regulates timely expression of a genetic network that contributes to early development across animals.

## RESULTS

### CPEB Phylogeny

We used the RBD domain of human CPEB1 (NP_001275748.1; AA 234-479) as input in TBLASTN searches (e-value cutoff = 0.05) against gene models of the following animal species: *Amphimedon queenslandica* (Porifera), *Mnemiopsis leidyi* (Ctenophora), *Nematostella vectensis* (Cnidaria), *Trichoplax adhaerens* (Placozoa), *Drosophila melanogaster* (Arthropoda), *Capitella teleta* (Annelida), *Schmidtea mediterranea* (Platyhelminthes), as well as *Mus musculus* and *Danio rerio* (Chordata). We also used the same query to search gene models from the following non-animal relatives: *Saccharomyces cerevisiae* and *Schizosaccharomyces pombe* (Fungi), *Arabidopsis thaliana* (Plantae), *Capsaspora ocwazarki* (Filasterea), and *Salpingoeca rosetta* (Choanoflagellatea). We detected sequences with RBD domains from all species except *Capsaspora ocwazarki* under the used parameters. After condensing genes with multiple isoforms down to one representative, we constructed an alignment of individual RRMs (Supplementary File S1) and used it to infer the phylogenetic relationships among individual RRMs present in CPEBs and related non-CPEB sequences. The RRMs of previously characterized CPEBs from *Drosophila*, the planarian *S. mediterranea*, and vertebrate species clustered together with hits that were exclusively from animal lineages (Figure 1C; Supplementary Figure S1). Sequences from non-animal lineages, including plant, fungi, and the choanoflagellate *S. rosetta*, were absent from clades that included RRMs from CPEB orthologs (Figure 1C; Supplementary Figure S1). Our results indicate that CPEBs are found in every major animal lineage and are specific to animals.

In addition to finding CPEBs in every major animal lineage, at least one ortholog each of CPEB1 and CPEB2 were identified in every metazoan species that we surveyed including Ctenophora and Porifera. This indicates that both CPEB1 and CPEB2 subfamilies were present in the genome of the last common ancestor of extant metazoans and suggests that the presence of both CPEB1 and CPEB2 is critical for most if not all animals. Notably, sequences corresponding to the first RRM (RRM1) of both CPEB1 and CPEB2 proteins clustered separately from those corresponding to the second RRM (RRM2) in all CPEBs (Figure 1C), indicating that CPEB1 and CPEB2 subfamilies share a common origin and conserve an ancestral architecture with tandem RRMs. Our parallel analysis using the Zinc Finger domain present in CPEBs (Supplementary Figure S2) was congruent with our analyses of the RRM domains.

The closest neighboring clade of RRMs to those of CPEBs contains factors known to be involved in post-transcriptional regulation of maternal mRNAs in bilaterians, including Squid and Musashi, DAZ Associated Protein-1 (DAZAP-1), and Heterogeneous nuclear ribonucleoprotein 27C (Hrb27C) orthologs. Musashi cooperates with CPEB in regulating cytoplasmic polyadenylation activity during vertebrate oocyte maturation [81–83], whereas DAZAP-1 binds to DAZL [84], which also cooperates with CPEB during oocyte maturation [85]. This clade also includes proteins from yeast, plant, and non-animal species closely related to metazoans, such as HRP1/YOL123W (which is required for 3’-end formation in *S. cerevisiae;* [86]), and heterogeneous nuclear ribonucleoproteins (hnRNPs) that participate in pre mRNA splicing, mRNA transport, and RNA editing [87, 88]. These proteins share a similar architecture as CPEBs in that they contain two RRMs in tandem, although these are positioned near the N-terminus rather than the C-terminus and lack a Zinc Finger (Figure 1D). As with CPEBs, members of this clade have RRMs that cluster into separate subclades (RRM1 and RRM2; Figure 1C). However, unlike CPEBs, these subclades were composed of sequences from different kingdoms, which suggests that these proteins were present in the last common eukaryotic ancestor. The third and last major clade found in our analysis included RRMs from orthologs of Nucleolin, which is a major regulator of rRNA biogenesis conserved in plants, yeast, and animals (reviewed by [89]), as well as splicing factors RSZ21 and RSZ22 from *A. thaliana* (Figure 1C). Altogether, these results support the hypothesis that CPEBs are a family of proteins exclusively present in animals. Additionally, these results suggest that CPEBs share ancestry with Musashi and DAZAP, and ultimately arose from heteronuclear RNA-binding proteins ubiquitously present in eukaryotes.

### Analysis of maternal mRNA regulation by cytoplasmic polyadenylation in Cnidaria

To test for evidence of cytoplasmic polyadenylation in cnidarian species, we extracted RNA from the ovaries, spawned eggs, embryos, and adult polyps of the sea anemone *Nematostella vectensis* at different timepoints post-fertilization and determined the expression of core cytoplasmic polyadenylation machinery components using reverse transcription PCR (RT-PCR; Figure 2, A-J). Expression of CPEB1 and GLD2 orthologs was readily detectable in the oocyte, the egg, and immediately post-fertilization, but declined at later stages of development (Figure 2J). Expression of the CPEB2 ortholog and a second GLD2 homolog (*NvGLD2-like*) were most obvious in samples obtained 10 and 40-days post-fertilization (dpf), which correlated the timing of expression of the neuronal and neuronal ectoderm marker *NvZicD* (Figure 2J; [90]). Expression of CPSF subunits (*NvCPSF73* and *NvCPSF100*) was detected at every tested time-point, as would be expected of factors that are involved with canonical 3’-end processing and polyadenylation in the nucleus (Figure 2J). In the absence of available antibodies to determine the presence of factors at the protein level, these results show expression of the cytoplasmic polyadenylation machinery in late oogenesis and early embryogenesis of *N. vectensis*, indicating that this is indeed a conserved feature between cnidarians and bilaterians.

**Figure 2.**
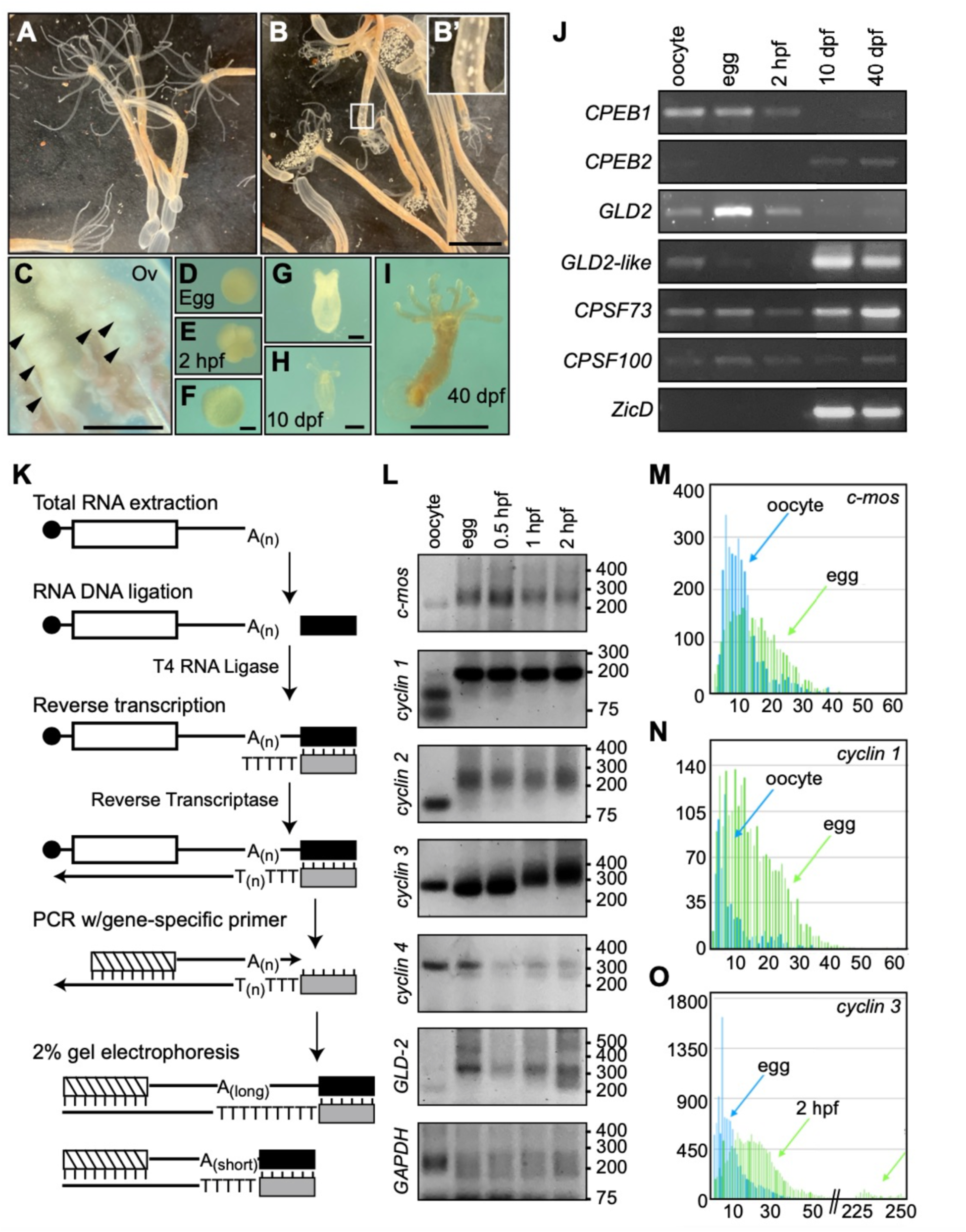
Cytoplasmic polyadenylation during oocyte maturation and early development of *Nematostella vectensis*. **(A-I)** Bright field images of *N. vectensis* reproductive process. Females before (A) and after spawning (B). Inset shows migration of eggs through column during ovulation (B’). Magnified view of ovaries (Ov; C), an egg (D), early cleavage (E), gastrula (F), tentacle bud stage (G), juvenile polyp (unfed; H), and polyp after multiple feedings (I). hpf, approximate hours post-fertilization at room temperature; dpf, days post-fertilization. Scale bars = 1 cm, B; 1 mm, C and I; 0.1 mm, F-H. **(J)** Developmentally regulated expression of cytoplasmic polyadenylation components detected by reverse transcription PCR. **(K)** PCR-based assay for measurement of poly(A) tail length. Total RNA extracted from specific tissues and developmental stages was ligated to the 5’-end of a DNA oligo and used as a template for reverse transcription using a primer annealed to the ligated DNA oligo. The synthesized cDNA containing the poly(A) region was then used as template to amplify the 3’-end of mRNAs of interest using a gene-specific primer and the primer antisense to the ligated DNA oligo. Changes in poly(A) tail length were assessed by differences in electrophoretic mobility of PCR products in a 2% agarose gel. See methods section for more details. **(L)** Assessment of shifts in electrophoretic mobility representative of changes in poly(A) length for gene specific transcripts using the assay shown in (K). Size of DNA markers is shown on the right. **(M-O)** Bar graphs depicting number of Amplicon-EZ reads (*x-*axis) for each specific length of poly(A) tail (*y-*axis) in *c*-*mos* (M), *cyclin 1* (N), and *cyclin 3* (O) mRNAs. Reads from oocytes are shown in blue and eggs in green in (M and N), whereas reads from eggs are shown in blue and those from embryos 2-hpf are shown in green in (O).

To determine whether long poly(A) tails result in increased translation products in *N. vectensis* eggs and embryos, we utilized a dual NanoLuc/Firefly Luciferase reporter system [91]. We used *in vitro* transcription to generate NanoLuc mRNAs with and without poly(A) tails of ∼100 to ∼350 adenosines in length, which is close to the initial length of ∼250 nucleotides observed in mammalian cells [92], and injected these into *N. vectensis* eggs and embryos (Supplementary Figure S3). We generated luciferase mRNAs lacking a poly(A) tail in the same manner and co-injected these as loading controls. Upon six hours post-injection, levels of NanoLuc and Luciferase activity indicated that polyadenylated mRNAs generated up to 30-fold more product than those lacking a poly(A) tail in both eggs and embryos (Supplementary Figure S3). To determine whether increased NanoLuc signal from polyadenylated reporters was due to higher mRNA stability, higher translation, or both, we compared the abundance of NanoLuc mRNA by reverse transcription quantitative PCR (RT-qPCR) in *Nematostella* eggs injected six hours prior. We observed that absence of a poly(A)-tail did not result in decreased stability of reporter mRNAs under these conditions (Supplementary Figure S3C). However, NanoLuc activity measured six hours post-injection in the same batch of injected eggs was over 10-fold higher for polyadenylated mRNAs than for counterparts lacking a poly(A) tail (Supplementary Figure S3, E-F). These data show that the presence of a poly(A) tail stimulates translation in *N. vectensis* eggs and embryos, as is known to occur during early development of bilaterians [93–96].

Next, we looked for evidence of poly(A) tail lengthening during ovulation and early embryonic development of *N. vectensis*. To do this, we identified *N. vectensis* homologs of known CPEB substrates, and measured poly(A) tail-length of their mRNA using a well-established PCR-based assay (Figure 2K; [97, 98]). Briefly, total RNA was extracted from dissected oocytes, eggs, and embryos at 0.5, 1, and 2 hours post-fertilization (hpf), and ligated with a DNA oligo at the 3’-end of the RNA. Then, a complementary oligo with a dT(5) extension at its 3’-end was used in a reverse transcription reaction selective for polyadenylated transcripts. The resulting cDNA was used as template for PCR using a gene-specific forward primer and a reverse primer identical to the one used for reverse transcription. Because the reverse primer anneals to the oligo originally ligated at the end of the transcripts, the presence of longer poly(A) tails results in upward shifts in electrophoretic motility when compared to shorter poly(A) tails on otherwise identical mRNAs (Figure 2K). Using this approach, we detected mRNA tail elongation for five conserved CPEB substrates (Figure 2L). Homologs of cell cycle regulators *c-mos (Nv_c-mos)* and *cyclin A* and *B* homologs (*Nv_cyclin 1* and *Nv_cyclin 2*, respectively) displayed longer poly(A) tails in the egg than in the oocyte (Figure 2L), matching what is known to occur during oocyte meiotic maturation in *Xenopus* [94]. The maternally expressed cytoplasmic poly(A) polymerase ortholog, *NvGLD2*, likewise displayed an increase in poly(A) tail length during meiotic maturation (Figure 2L), which is also observed in *Xenopus* [97]. A third cyclin homolog, *Nv_cyclin 3* (closest human homolog being cyclin B3), was polyadenylated upon fertilization, most noticeably between 0.5 and 1-hpf (Figure 2L). Conversely, the poly(A) tail of mRNAs encoding for a fourth cyclin homolog (*Nv_cyclin 4*), as well as the housekeeping gene *GAPDH*, either stayed constant or shortened as development progressed (Figure 2L). We verified that the electrophoretic motility of amplicons from polyadenylated transcripts matched the size predicted based on the position of the gene-specific primer, the length of 3’ends (according to 3’UTR reads deposited at the Stowers Institute for Medical Research repository (simrbase.stowers.org) and the National Center for Biotechnology Information (NCBI; www.ncbi.nlm.nih.gov), plus potential poly(A) tails), and the ligated 3’end adapter (Table 1; Supplementary Figure S4). This process also revealed the presence of canonical CPEs in the 3’UTR of each of the polyadenylated mRNAs (Supplementary Figure S4). Altogether, these results show dynamic lengthening and shortening of maternal mRNAs of *Nematostella* in a manner that parallels behaviors observed during early development of bilaterians. The increases of poly(A) tail length observed in *N. vectensis c-mos*, *GLD-2*, and *cyclin 1*, *2*, and *3* homologs, as well as the presence of CPEs in their 3’UTRs, support the hypothesis that CPEB-mediated polyadenylation during meiotic maturation and early development are part of a conserved genetic program that predates the last common ancestor of cnidarians and bilaterians.

**Table 1.**
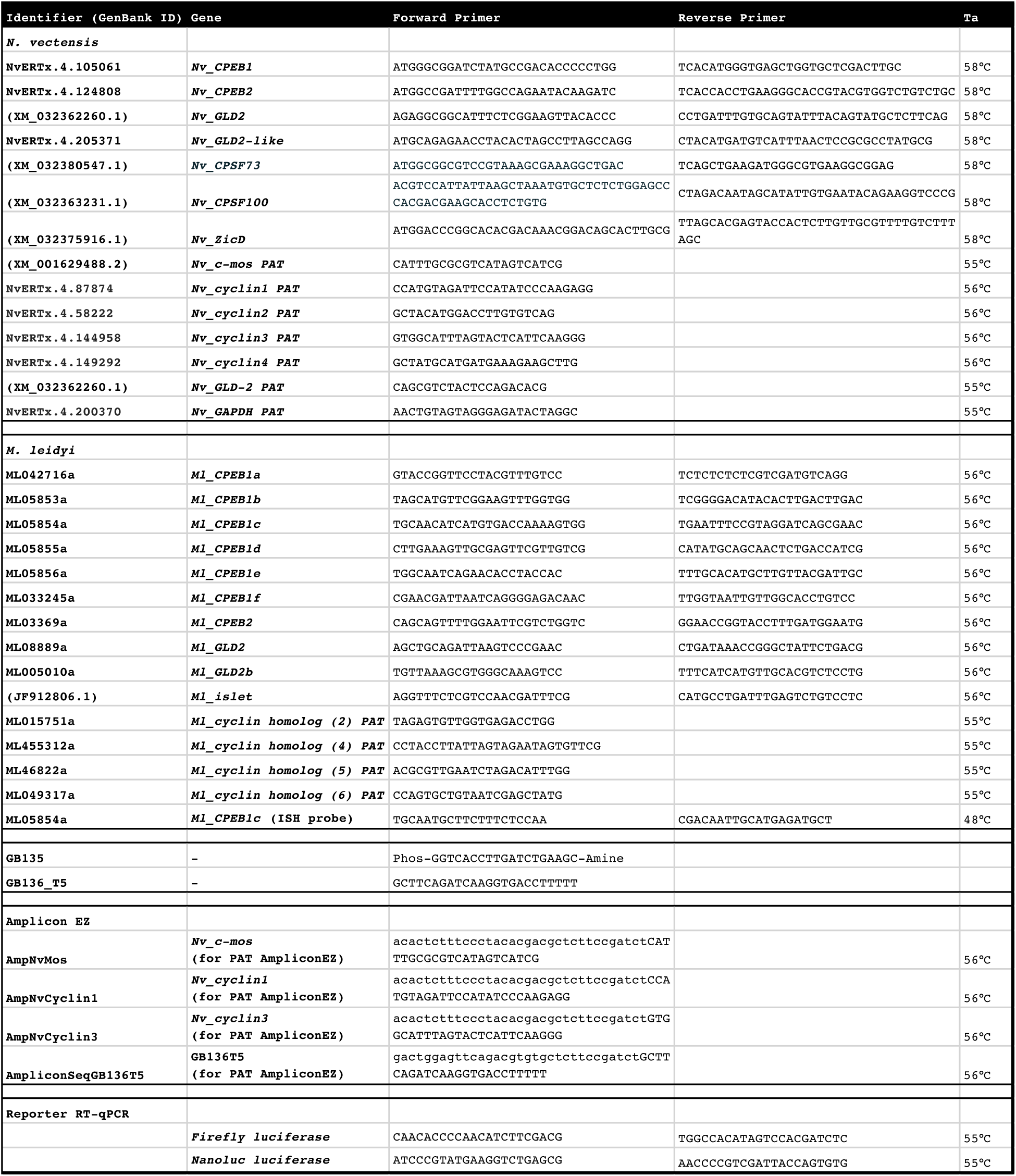
Oligonucleotides used in this study. The sequence of oligonucleotides used as primers for gene specific amplification of cDNA (Forward and Reverse), as well as poly(A) tail length assays (PAT) and reverse transcription-quantitative PCR (RT-qPCR), are listed along with corresponding annealing temperatures (Ta).

We sought to validate the identity of amplicons produced in our poly(A) tail length assay directly by Sanger sequencing fragments cloned into bacterial vectors. During this process, we verified the identity of our PCR products and presence of poly(A) tails of different lengths (Supplementary Figure S4 and S5). We also observed evidence of non-templated incorporation of uridine and guanosines within the poly(A) of some transcripts (Supplementary Figure S5). To investigate the changes in length and composition of poly(A) tails represented in our PCR-based assay in more depth, we analyzed hundreds of amplicon sequences for *c-mos*, *cyclin 1*, and *cyclin 3* mRNAs using Illumina Next-Generation sequencing of PCR amplicons (Amplicon-EZ, Genewiz, Azenta Life Sciences, South Plainfield, NJ; Supplementary Figure S6). Cognizant of the caveats that sequencing through long stretches of polynucleotide repeats is difficult, and that measurements using sequencing approaches tend to underestimate the precise length of poly(A) tails [99, 100], we designed a program that selects for tail sequence represented in both forward and reverse Amplicon-EZ reads (get_polya.pl, in GitHub repository). This program trims genic sequence and potential sequencing errors and calculates the number of non-templated positions at the 3’-end of mRNAs represented in our Amplicon-EZ reads (Supplementary File S2). Using this approach, we detected extension of poly(A) tail length during oocyte maturation for *c-mos* and *cyclin 1*, as well as after fertilization for *cyclin 3*. For *c-mos,* we detected an increase in poly(A) tail length from a mean of 10 nucleotides in oocytes to 15 nucleotides using this method (median 9 and 13 nucleotides, respectively; Figure 2M, Supplementary Figure S6). Likewise, we detected an increase in mean from 8 to 14 nucleotides for *cyclin 1* (median of 5 to 12 nucleotides, respectively; Figure 2N, Supplementary Figure S6) and from 9 nucleotides in the egg to 38 nucleotides in embryos 2-hpf for *cyclin 3* (median 7 and 20 nucleotides; Figure 2O, Supplementary Figure S6). Interestingly, two peaks of poly(A) tail length (at median 19 and 247 nucleotides) were observed when plotting the length of *cyclin 3* poly(A) tail at the latter developmental time-point (2-hpf; Figure 2O), suggesting a potential heterogenous population of cytoplasmic polyadenylation products from the same gene. Overall, we observed a trend of poly(A) tail lengthening in developmental timepoints that correlates with those observed by gel electrophoresis for all three genes (Figure 2L), indicating that the shifts in electrophoretic motility of amplicons observed in our original PCR-based poly(A) tail assays are due to changes in poly(A) tail length.

Analysis of nucleotide composition of poly(A) tails from Amplicon-EZ data revealed that adenines were by far the most prevalent base, encompassing ∼76% of all positions in the tail (Supplementary Table S2). Uridine was the second highest by occupying ∼16% of all positions. Guanines and cytosines composed ∼5% and ∼3%, respectively. We observed differences in composition between the 5’ and 3’ regions of the tail, with uridine enrichment at immediate positions after the cleavage and polyadenylation site (Supplementary Table S2). To visualize the mixed composition of the 5’-end of these tails we depicted the frequency of each nucleotide in the first 20 positions of untemplated sequence for tails of at least 5 nts using the WebLogo sequence generator [101]. Using this approach, we observed similar occupancy of uridines and adenines at the first position of the tail, with gradual decreases in prevalence of uridine moving downstream (Supplementary Figure S6). Because the high presence uridine at initial positions of the tail was not as prevalent in sequenced cDNA clones (Supplementary Figure S5), we performed additional tail composition analysis using of *N. vectensis* egg RNA using Illumina RNAseq reads from ribodepleted samples (Supplementary Figure S7), as well as Nanopore cDNA sequencing of total RNA (Supplementary Figure S8). We found that reads representing *c-mos*, *cyclin1*, and *cyclin 2* mRNAs in Illumina reads included uridines within their poly(A) tails (Supplementary Figure S7). However, the presence of uridine in the first or second position of the tail in Illumina reads was only observed once each in >30 reads. Furthermore, uridines were absent in all the poly(A) tails corresponding to *cyclin 3* mRNA (Supplementary Figure S7), suggesting that incorporation of oligouridine may not as prevalent as represented in Amplicon-EZ reads. Sequences obtained using Nanopore technology had adenosines as the most prevalent base, but uridines were also present in most tails of analyzed mRNAs (12/13 for *c-mos*; 21/32 for *cyclin1*, and 30/38 for *cyclin3*; Supplementary Figure S8). However, the of uridines were throughout the tail and not enriched at either end (Supplementary Figure S8). While absence of correlation regarding the distribution of non-adenosine bases within the tail of mRNAs prohibits conclusive interpretation of these data, the frequent observation of mixed tails using four different approaches merits future investigation. Nevertheless, our data using RNA extracted from different developmental stages analyzed in parallel by electrophoretic mobility and AmpliconEZ sequencing clearly indicate that poly(A) tail lengthening for *cyclin* and *c-mos* mRNAs takes place during oogenesis and early development in *N. vectensis*.

### Analysis of cytoplasmic polyadenylation in ctenophores

Our phylogenetic analyses revealed the existence of CPEB1 and CPEB2 orthologs in the genome of the ctenophore *Mnemiopsis leidyi* (Figure 1C; Supplementary Figures S1 and S2). We identified six *M. leidyi* genes grouped as members of CPEB1 subfamily (Figure 1C; Supplementary Figure S1 and S2), which was surprising because the CPEB1 family is most often represented by a single gene in previously analyzed species. To determine where within Ctenophora the CPEB1 expansion occurred, we looked for CPEB homologs in the genomes of *Hormiphora californensis* [102] and *Beroe ovata* (Hernandez, Ryan, *et al., unpublished*). In both of these ctenophore genomes, we identified one CPEB2 and six CPEB1 sequences (Supplementary Table S3; Supplementary Figure S9). These results suggest that the expansion of CPEB1 occurred in the last common ancestor of these three ctenophores and has been maintained in all three lineages.

We asked whether maternal expression of the CPEB1 ortholog and components of the cytoplasmic polyadenylation complex was a conserved feature in *M. leidyi*. Most ctenophores, including *M. leidyi*, are hermaphrodites that develop testes and ovaries parallel to each other under each of eight comb-rows that are aligned along the oral-aboral axis (Figure 3, A-D). The presence of mesoglea and proximity of comb plates make it difficult to specifically isolate oocytes or ovarian tissue (Figure 3, A and B). In addition, *M. leidyi* is able to self-fertilize very effectively [103] prohibiting analysis of unfertilized eggs. However, hundreds of synchronized embryos can be obtained following controlled spawning events in the laboratory (Figure 3E; [104, 105]). Upon spawning, cleavage occurs every 15–20 minutes (Figure 3, F-K) and gastrulation can be observed by 4-hpf (Figure 3L; [103]). The juvenile cydippid stage is achieved by 20-hpf (Figure 3M), at which point specimens can eat, grow, and eventually develop gametes.

**Figure 3.**
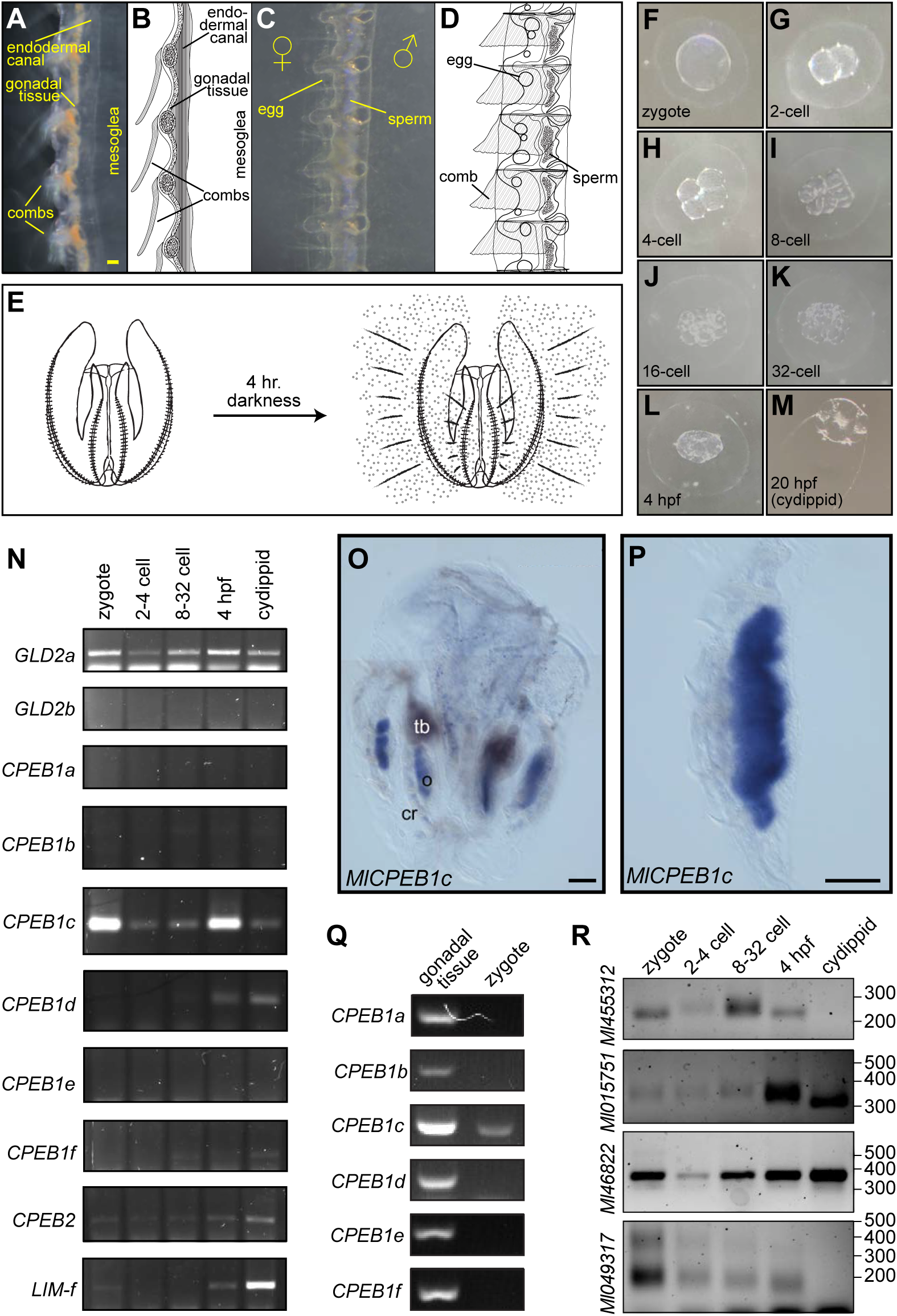
Cytoplasmic polyadenylation in the ctenophore *Mnemiopsis leidyi*. **(A-E)** Anatomy of *M. leidyi* reproductive structures and induced spawning. Dark-field microscopy images (A, C) and corresponding diagrams (B, D) depicting sagittal (A-B) and frontal (C-D) views of the gonadal structures positioned between the comb rows and mesoglea of *M. leidyi*. Male and female gametes are present on opposing sides along the midline of comb rows in lobate stage *M. leidyi*. (E) Illustration depicting induction of *M. leidyi* spawning in cultures maintained in the laboratory under constant light exposure by placing in the dark for 4 hrs. **(F-M)** Bright field images representative of developmental progression of *M. leidyi* embryos (hpf, hours post-fertilization). **(N)** Detection of developmentally regulated expression of cytoplasmic polyadenylation complex components GLD2 and CPEB, as well as the ectodermal marker *Lim-f*, by reverse transcription PCR. **(O-P)** Differential Interference Contrast microscopy image of *in situ* hybridization samples displaying *MlCPEB1c* mRNA expression in ovaries of a sexually mature cydippid (O) and in a dissected sample of a comb row with gonad (P). Abbreviations: ovary (o), comb row (cr), tentacle bulb (tb). **(Q)** Detection of expression of CPEB paralogs and the ectodermal marker *Lim-f* by reverse transcription PCR in lobate-stage ctenophore comb-rows containing gonadal tissue, zygotes, and cydippids. **(R)** PCR-based assessment of poly(A) tail length (as in Figure 2K) shows differences in 3’-end length for mRNAs of *M. leidyi* cyclin homologs at different developmental stages. Position of DNA size markers is shown on the right. Scale bars = 0.1 mm. Embryos displayed in (F-L) are approximately 0.2 mm in diameter.

We extracted RNA from embryos collected as zygotes, combined 2 to 4-cell stages, combined 8 to 32-cell stages, and 4-hpf, to assess expression of CPEBs by reverse transcription PCR. We also included RNA extracted from unfed cydippids in our analyses as control for absence of maternal products and expression of markers for differentiated tissue. Using gene-specific primers, we detected maternal expression of one CPEB1 paralog (*MlCPEB1c*; identifier ML05854a in the *Mnemiopsis* Genome Project Portal [106]) and one of two GLD2 cytoplasmic poly(A) polymerase homologs (*MlGLD2a*; ML0889a) in zygotes (Figure 3N). Expression of four of the six CPEB1 paralogs (*MlCPEB1a*/ML042716; *MlCPEB1b*/ML05853a; *MlCPEB1e*/ML05856a; and *MlCPEB1f*/ML033245a) was not conclusively detected at any stage, nor was expression of a second GLD2 homolog (*MlGLD2b*/ML005010a). *MlCPEB1d* (ML05855a) and the CPEB2 ortholog (*MlCPEB2*/ML03369a) were both detected at the 4-hpf time-point and in cydippids, but not during earlier stages of development (Figure 3N). The timing of expression of *MlCPEB1d* and *MlGLD2b* mimic the expression of *Ml_islet* (ML053012a), a LIM family gene expressed in ectoderm and the apical organ of cydippids [107]. These results indicate that at least one member of the CPEB1 family and one GLD2 poly(A) polymerase are maternally expressed in *M. leidyi*.

We sought to determine whether *MlCPEB1c* is expressed maternally by direct analysis on developing oocytes. To do so, we obtained sexually mature *M. leidyi* cydippids that had visible gonads and determined the distribution of expression of *MlCPEB1c* by whole-mount *in situ* hybridization (Figure 3, O and P). Using this approach, we observed robust expression of *MlCPEB1c* in developing oocytes (Figure 3, O and P). Detection of *MlCPEB1c* expression was restricted to the ovary, which shows parallels to what is observed in other animals [38, 43]. Other CPEB1 paralogs were not detected in zygotes (Figure 3N) but were detected by reverse transcription-PCR in RNA extracted from dissected comb-rows containing gonadal tissue from lobate-stage ctenophores (Figure 3Q). Altogether, these results indicate that maternal expression of CPEB1 orthologs is an ancestral feature that is shared across metazoan lineages, which is consistent with the notion that regulation of gene expression by cytoplasmic polyadenylation is essential for early development in animals. *MlCPEB1c* is available in the zygote to regulate maternal mRNAs in *M. leidyi*, while other CPEB1 orthologs may function at different stages of germline (sperm or egg) development.

Given the evidence for maternal deposition of cytoplasmic polyadenylation machinery in ctenophores, we asked whether conserved targets display changes representative of poly(A)-tail elongation during early development of *M. leidyi*. To do this, we identified homologs of cell cycle regulators present in the *M. leidyi* transcriptome using TBLASTN and designed gene-specific primers close to the end of their 3’-UTR for analysis of poly(A)-tail length changes as in *N. vectensis* (Figure 2K). Using this approach, we looked for changes in electrophoretic mobility of 3’-end amplicons for four *cyclin* homologs during the earliest stages of development (Figure 3R). Amplicons of one *cyclin* homolog (identifier ML455312a) became longer during the transition from zygote to the 32-cell stage, decreased in size at 4-hpf, and were absent in cydippids 20-hpf (Figure 3R), indicating potential poly(A)-tail lengthening during the first embryonic cleavage events, followed by deadenylation and decay at later stages of development. Two other *cyclin* homologs displayed shortening as development progressed (Figure 3R). One of these (ML015751a) remained relatively unchanged throughout the first developmental stages analyzed, but decreased in size in cydippids, while the other (ML049317a) was shortened gradually and was undetectable in cydippids (Figure 3R). The size of amplicons from one of these *cyclin* homologs remained unchanged throughout the analysis (ML46822a). Intrigued by the potential changes in poly(A) tail length observed for ML455312a during the initial embryonic cleavages, we performed a more detailed assessment of poly(A) tail length changes that included analysis of each of the first four cleavages separately (Supplementary Figure S10). This approach revealed alternating increases and decreases in amplicon size following each of the first three divisions (Supplementary Figure S10). To verify the identity of our amplicons, we cloned and sequenced cDNA from the zygote and two-cell embryos, which validated the genetic identity of ML455312a PCR products and revealed changes in poly(A) tail composition from average of 15.7 in the zygote to ∼32 adenosines in two-cell embryos (Supplementary Figure S5). We conclude that cytoplasmic polyadenylation of maternal transcripts takes place during the initial cleavages of *M. leidyi* embryos, and that cytoplasmic polyadenylation of *cyclin* homologs is a conserved feature in ctenophores, cnidarians, and vertebrates. Altogether, our results support the hypothesis that an ancestral network of post-transcriptional regulation via cytoplasmic polyadenylation is a ubiquitous feature of early development amongst animals.

## DISCUSSION

In 1983, Evans et al. [108] reported the identification of cyclins as proteins made from maternal mRNA and destroyed after cell division. Years of research have uncovered how robust transcriptional regulation of cyclin genes drives progression between stages of the cell cycle in somatic cells across Eukarya [109, 110]. Here, we provide evidence indicating that regulation of cyclins via cytoplasmic polyadenylation is a unifying theme in animal development. We postulate that generation of cyclins from a pool of stored mRNAs was an important innovation for evolution of multicellular development. Post transcriptional regulation by cytoplasmic polyadenylation allows cells to bypass the requirement of generating new transcripts, which would impede the quick S to M phase transitions observed in early embryonic development. In absence of the ability to turn maternal mRNAs off and on, rapid mitotic divisions would have to pause between stages to allow for novel transcription to occur, bringing along large changes in gene activation programs. The enzymatic extension of poly(A) tails provides a mechanism for tunable and sequential activation of different groups of maternal mRNAs, as observed during progression from oocyte to egg, to embryo [60, 83, 94, 111].

### Origin of CPEBs

This study reveals the presence of CPEB1 and CPEB2 orthologs in every major animal lineage and provides evidence for cytoplasmic polyadenylation of maternal mRNAs in ctenophores and cnidarians. While genetic and biochemical perturbations are necessary to conclusively determine the involvement of CPEB in cytoplasmic polyadenylation of maternal mRNAs in *M. leidyi* and *N. vectensis*, the timing of detection of cytoplasmic polyadenylation machinery and poly(A) extension of conserved targets during early development, strongly parallels what is known about function of CPEB in bilaterians. Unlike other essential components of the cytoplasmic polyadenylation machinery, such as non-canonical poly(A) polymerases of the GLD2 family [77, 78, 112] and subunits of the Cleavage and Polyadenylation Specificity Factor, clear CPEB orthologs are only present in metazoans. Indeed, we did not find CPEBs in Filasterea or in choanoflagellates, the latter of which are by many measures the closest relatives to animals [113, 114]. Given their absence in non-animal life forms, as well as their conserved function during the transition from oocyte to zygote to multicellular embryo, we postulate that emergence of CPEBs may have been a pivotal factor in evolution of multicellular development from single celled animal ancestors.

Interestingly, the closest CPEB relatives in animals include post-transcriptional regulators of meiosis and early development, such as Musashi, Squid, Hrb27C, and DAZ-Associated Protein-1 (DAZAP1). Both Musashi and DAZL work with CPEB to coordinate timely translation of maternal mRNAs in vertebrates [81–83, 85], whereas Squid and Hrb27C cooperate to regulate mRNA localization and translation in *Drosophila* oocytes [115, 116]. These RNA-binding proteins may have coevolved to modulate the identity of their targets, as well as the timing and strength of translational activation/repression in specific animal lineages, while maintaining a “core” network of shared targets. Our phylogenetic analysis of RRMs grouped the closest non-animal relatives of CPEB with HnRNPs present in fungi, plants, and choanoflagellates (Figure 1C). The functions of many of these proteins remains to be characterized, but the yeast protein that is most similar to CPEBs (Hrp1) is known to participate in transcriptional termination and 3’-end processing, as well as to shuttle between the nucleus and the cytoplasm, and mark messages for nonsense-mediated decay [86, 117, 118]. Characterization of more members of this group of RRM-containing proteins will be needed to form a formal hypothesis regarding the origin of CPEBs. One possibility is that that CPEBs derived from ancestral nuclear HnRNPs that participate in 3’-end processing and maintained interactions with CPSF.

### Ancestral CPEB *architecture*

High primary sequence conservation is found in the C-terminal region across CPEB orthologs (40% to 90%) and paralogs (> 30%), with highest identity being shared amongst CPEB2 subfamily members [38, 119]. This can be predicted to reflect not only strong selection for the specificity of RNA binding provided by RRMs that reside in this region of the protein, but also of its participation in conserved protein-protein interactions. For example, the Zinc-Finger domain at the end of the RBD of CPEBs is predicted to bind sumoylated proteins required for cytoplasmic polyadenylation, such as CPSF and the scaffolding protein Symplekin [31, 120]. Surprisingly, not all CPEB partners interact with the conserved region of the protein. For example, members of the PUF family of proteins work with CPEBs to maintain repression of target mRNAs in nematodes, flies, and vertebrates [121, 122]. Physical association between the CPEB protein CPB-1 and the PUF family member FBF-1 from *Caenorhabditis elegans* was mapped to a small motif that resides in a disordered region upstream of the RBD [123]. In addition, motifs for phosphorylation and ubiquitination of CPEB, which modulate regulated timing of activation of cytoplasmic polyadenylation and destruction of CPEB (respectively), also reside outside of the conserved RBD [124–127]. Nevertheless, sequence conservation outside of the RBD of CPEBs from different phyla is not prevalent, and this may reflect highly specific needs for regulation of CPEB activity in each species or limitations in our programs for sequence analysis.

One domain that has been characterized in the N-terminal region of some CPEBs are stretches of glutamine-rich sequence that mediate prion-like conformational changes [52, 119, 128, 129]. Although their position and presence across orthologs are not conserved, stretches of polyglutamine in CPEB orthologs from *Drosophila* and the marine snail *Aplysia* are required for proper memory formation [128, 130, 131] reviewed by [132]. Whereas the *Aplysia* protein originally found to form prions is a CPEB1 ortholog, prion domains are found in CPEB2 subfamily members in mammals, planarians, and flies. Amongst ctenophores, putative prion domains are present in *M. leidyi* CPEB1a and a CPEB1 paralog in *Beroe ovata* (Supplementary Table S3). Furthermore, we identified putative prion domains in both CPEB1 and CPEB2 orthologs in the ctenophore *Hormiphora californensis* (Supplementary Table S3). The presence of prion domains in CPEBs across major animal lineages suggests that prion-mediated aggregation may be a shared characteristic that facilitates some contribution(s) that cytoplasmic polyadenylation brings to animal biology. However, the distribution of prion domains suggests that these may be products of convergent evolution. It will be interesting to see whether some CPEBs utilize prion conformation to form aggregates of ribonucleoprotein in the germline, as thus far is known to occur in neurons.

### Correlation between length of poly(A) tail and mRNA stability or translation

The poly(A) tail of an mRNA protects the 3’-end of the message from degradation and promotes translational efficiency [133, 134]. Benefits bestowed by the poly(A) tail are mediated by cytoplasmic poly(A) binding proteins (PABPCs), which bind to poly(A) every 27 residues upon mRNA export into the cytoplasm [22, 135–137]. PABPCs can also bind translation initiation factors attached to the 5’-cap, forming a bridge between both ends of the message and arranging mRNAs into “closed-loop” structures that are believed to support the stability and recycling of ribosomes [137, 138]. Poly(A)-binding proteins can also directly stimulate translational initiation *in trans* [139], or when tethered to a message [140, 141], regardless of the presence of a poly(A) tail.

Recent studies have shown that the correlation between poly(A) tail length and translational efficiency is absent in somatic cells and lost after zygotic genome activation [142–144]. One explanation for the uncoupling between poly(A) tail length and translational efficiency is that levels of PABPCs are rate-limiting in the oocyte but increase later in development [145]. An alternative hypothesis, although not mutually exclusive from the one aforementioned, is that multiple units of PABPC must be bound to an mRNA for translational stimulation. A minimum of 12 adenosines is required for Poly(A) Binding Protein and multiples of 27 for assembly of multiple units [135, 146, 147]. However, this second hypothesis is challenged by the observation that the correlation between poly(A) tail length and translational efficiency is lost after the ∼20 nucleotide threshold in HeLa cells, which suggests that binding by one PABPC may be enough for translational stimulation [143], as well as by the observation that repressed *c-mos* and *cyclinB1* mRNAs have an average length of 50 and 30 adenosines (respectively) when repressed in *Xenopus* oocytes [94]. A third hypothesis posits that the type and combinations of PABPCs in specific cell types lead to different dynamics between poly(A) tail length and translational efficiency [148].

While further studies are needed to measure the precise length of poly(A) tails in ctenophores and cnidarians, it is worth contemplating the observation that tails of repressed cytoplasmic polyadenylation targets in *N. vectensis* may be shorter than the minimal requirement for binding a single PABPC (Figure 2, M-O; Supplementary File S2; Supplementary Figure S4). This observation suggests that either PABPC binding is dispensable for protection from mRNA decay in oocytes and eggs of *N. vectensis*, or that PABPC binding has a smaller footprint in these species. Indeed, NanoLuc reporter mRNAs lacking a poly(A) tail were as stable as polyadenylated counterparts in injected *N. vectensis* eggs (Supplementary Figure S3D). Conversely, the polyadenylated status for conserved CPEB substrates in *Nematostella* yielded tails that sometimes surpassed the 100-nucleotide mark (Figure 2, L and O), whereas lengthening of the poly(A) tail in *Mnemiopsis* were much more modest (albeit sampling the latter is limited to one cyclin gene; Figure 3R and Supplementary Figure S10). Future global surveys of poly(A) dynamics during development will lead to an understanding of whether shorter poly(A) tails are the norm in *M. leidyi*, and the degree to which translational enhancement is provided by small increments (20-30 nucleotides) in poly(A) tail length in ctenophores and cnidarians.

### Composition of mRNA tails

**Figure 4.**
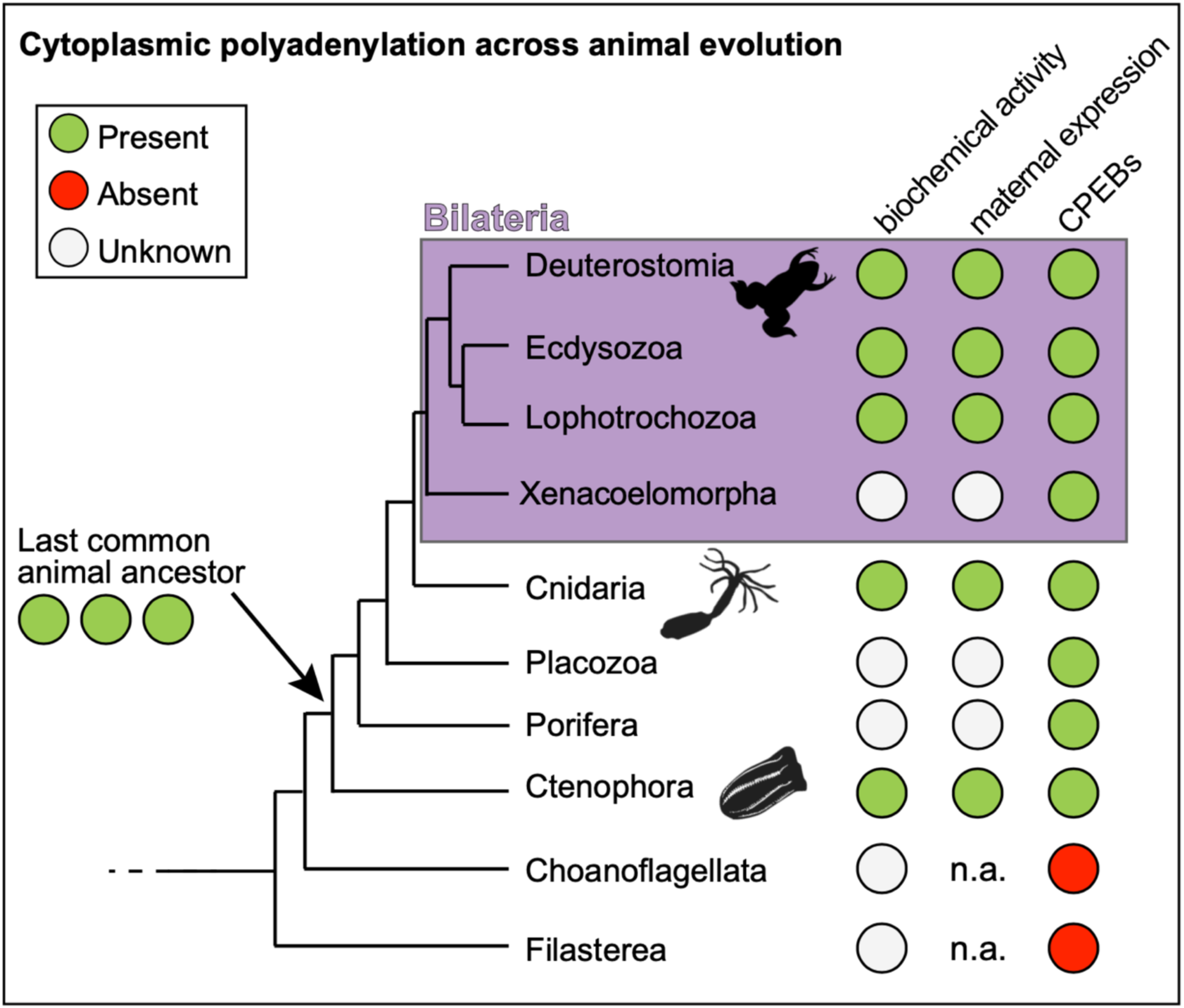
Working Model. Incorporation of results from this study with studies on bilaterians (magenta) demonstrate evidence for the presence (green circles) of CPEBs (right column) and their maternal expression (middle column) across animals. This work also demonstrates cytoplasmic polyadenylation of conserved targets across animal evolution (left column). The absence (red circle) of CPEBs in genomes of choanoflagellates and Filasterea indicates that cytoplasmic polyadenylation arose in the stem lineage of animals. White circles indicate that information is not available for a particular lineage. Animal relationships are based on Ryan et al. (2013). Abbreviation: not applicable, n.a.

The GLD-2 family is composed of ribonucleotidyl transferases that synthesize poly(A), but it also includes poly(U) polymerases (a.k.a. TUTases), poly(A) polymerases that incorporate intermittent guanosines within the poly(A) tail, and even polymerases that generate tails of (GU) repeats [77]. While oligouridylation by TUTases marks RNAs for degradation [149–151], the description of mixed tailing in the literature remains relatively novel and scarce in comparison. Recently, detection of poly(UG) tails was reported in targets of RNA-interference and transposons in *C. elegans*, where they serve as template for RNA-dependent RNA polymerase synthesis of trans-generational siRNAs [152]. Intermittent incorporation of guanosine in the poly(A) tail has recently been shown to stall deadenylation machinery and prolong the life of specific maternal mRNAs in zebrafish and *Xenopus* embryos [153]. TENT4 proteins are the enzymes responsible for intermittent guanylation [153], but intermittent guanosines are also deposited by GLD2 (*i.e.*, TENT2) orthologs when tethered to reporter RNAs [112]. We observed intermittent guanosines in poly(A) tails of maternal mRNA from *N. vectensis*, suggesting that this feature of tail regulation may also be ancestral to bilaterians and cnidarians, and may result from enzymes with lower fidelity than nuclear Poly(A) Polymerases.

Uridylation at the 3’-end of short poly(A) tails has been observed in animals, plants, and fungi [154–156]. We observed instances of non-templated uridines at the start of some analyzed mRNA tails from *N. vectensis*. However, clear differences in prevalence and position of uridine within the tail were observed between sequencing approaches; Supplementary Figure S5-S8). When using Amplicon-EZ sequencing, oligouridine was observed at the first positions of the tail almost as often as adenine and decreasing in prevalence downstream (Supplementary Figure S6, C-E; Supplementary File S2). Similar observations were reported for *cyclinB* mRNA in starfish oocytes, where oligouridine present at the initial positions of the tail was slightly trimmed during meiotic progression and cytoplasmic polyadenylation [157]. Uridines at the start of the tail do not seem to be conducive to decay in the oocyte or the egg and remain present even after cytoplasmic polyadenylation [157]. Ochi and Chiba hypothesized that oligouridine at the start of the tail may be involved in translational inactivation of mRNAs. Another possibility is that runs of uridine present at the start of the tail serve as a pre-existing mark to accelerate decay, and these become functional after deadenylation and the activation of mRNA degradation machinery during the maternal-to-zygotic transition [158]. Further studies will be needed to determine the pervasiveness of this structure on maternal transcripts of different animal species, as well as the identity of the polymerase responsible for incorporating uridine at beginning positions of the tail.

## MATERIALS AND METHODS

### Reproducibility and transparency statement

Custom scripts, command lines, and data used in these analyses, including sequencing reads, as well as alignments and tree files, are available at https://github.com/lrouhana/cpeb_evolution. To maximize transparency and minimize confirmation bias, we planned analyses a priori using phylotocol [159] and posted this original document and any subsequent changes to our GitHub repository (URL above).

### Identification and phylogenetic analyses of CPEBs

We identified putative CPEBs and outgroup sequences with the following approach. We used BLASTP version 2.10.1 with the query sequence Human CPEB1 (NP_001275748) against a database that included protein models from the following species: *Saccharomyces cerevisiae*, *Schizosaccharomyces pombe*, *Arabidopsis thaliana*, *Capsaspora owczarzaki*, *Salpingoeca rosetta*, *Amphimedon queenslandica*, *Nematostella vectensis*, *Trichoplax adhaerens*, *Drosophila melanogaster*, *Capitella teleta*, *Mus musculus*, *Danio rerio*, and *Mnemiopsis leidyi.* We also ran the same BLASTP using the web interface at PlanMine version 3.0 [160] to identify CPEBs and related sequences from *Schmidtea mediterranea*.

We downloaded HMMs from PFAM: RRM_1 (PF00076), RRM_7 (PF16367), and CEBP_ZZ (PF16366). For CEBP_ZZ, which should only occur once per CPEB protein, we created an alignment of CEBP_ZZ domains using hmm2aln.pl version 0.05 (https://github.com/josephryan/hmm2aln.pl), which uses hmmsearch from HMMer version 3.3 [161] to identify target domains in protein sequences and construct an alignment to the query hidden Markov model. For the RRM_1 and RRM_7 HMMs, which often produce overlapping results, we ran hmmsearch separately with each of these HMMs and merged overlapping results by taking the lowest N-terminal coordinate and the highest C-terminal coordinate from the 2 results. We extracted the amino acid sequences between the start and end of the match and used MAFFT version 7.407 [162] with default parameters to generate an alignment of RNA recognition motifs (RRMs). We used IQ-TREE version 1.6.12 [163] with default parameters to generate a maximum likelihood tree from all resulting alignments. All command lines, scripts, and sequence sources used in this section are available in our GitHub repository (https://github.com/lrouhana/cpeb_evolution). We used a web interface to perform reciprocal best BLAST searches to identify CPEBs in *Beroe ovata* and *Hormiphora californensis* (see Supplemental Table S2).

### Husbandry and spawning of animals in the laboratory

Husbandry of laboratory lines of *Nematostella vectensis* was performed as per [164]. Briefly, separate *N. vectensis* male and female colonies were maintained in separate bowls at 17 °C in 1/3 seawater under dark conditions and fed freshly hatched *Artemia* 1-2 times per week. We induced spawning of sexually mature animals by overnight exposure increased temperature and light as described by [165] following ingestion of minced oyster 24-48 hours prior. Upon spawning, eggs were de-jellied using 3% cysteine solution in 1/3 seawater (pH ∼7.5), fertilized and/or injected with reporter mRNAs, and collected for RNA extraction in TRI Reagent (Sigma-Aldrich, St. Louis, MO). Injections and development of fertilized eggs were performed in 1/3 filtered seawater at room temperature as described by Leyden et al. [166].

We collected *Mnemiopsis leidyi* hermaphrodites from floating docks on waters surrounding the Whitney Laboratory of Marine Biosciences, Marineland, FL, and maintained them as per Ramon-Mateau et al. [167]. We induced spawning by interrupting continuous light exposure for ∼4 hrs. as per Pang and Martindale [104], then visualized embryos under light microscopy and collected at the desired stages manually to freeze immediately at -80 °C in TRI Reagent for RNA extraction.

### Generation of cDNA and assessment of gene expression by reverse transcription-PCR

Total RNA was extracted from *N. vectensis* ovaries, or groups of 20 eggs, embryos, and polyps using TRI Reagent as per manufacturer’s instructions (Sigma-Aldrich, St. Louis, MO) and resuspended in 8 μl of RNase-free water. One microgram of total RNA was then ligated to 0.4 μg of GB135 3’-amino modified DNA anchor primer (Table 1) as per [98, 168], using T4 RNA ligase (New England Bioloabs, Ipswich, MA) in 10 μl volume reactions. After 2-hour incubation at 25 °C, and heat-inactivation of T4 RNA Ligase through 15 minutes of incubation at 65 °C, the contents of the ligation reaction were used as input for reverse transcription. 50 μl SuperScript IV Reverse Transcription reactions were performed a per the manufacturer’s instructions (Invitrogen, Carlsbad, CA), using the entire volume of corresponding T4 RNA ligation reaction and 0.5 μg of GB136_T5 primer (Table 1). The reverse transcription reaction was heat inactivated by incubation at 85 °C for 5 minutes. Then, one microliter volumes of Reverse Transcription reactions were used as template for amplification of internal cDNA fragments using in 20 μl volume PCR reactions (35 rounds of amplification using Promega 2X PCR Master Mix (Promega, Madison, WI). Gene-specific primers and annealing temperatures are listed in Table 1. The identity of PCR products was verified by Sanger sequencing of amplicons cloned using the pGEM-T cloning system (Promega, Madison, WI) as per manufacturer’s instructions. New sequence records were deposited in NCBI under GenBank accession numbers OP806308-OP806311 and OP852652-OP852653.

*Mnemiopsis leidyi* cDNA synthesis was performed as described above, but RNA was extracted from groups of 50 embryos or cydippids and resuspended in 6 μl of RNase-free water. Five microliters from the total resuspension were used as input for ligation with GB135 oligo using T4 RNA-ligase. The following steps proceeded as detailed above using the primers and annealing temperatures listed in Table 1. For analysis of CPEB paralog expression in comb-rows containing gonadal tissue dissected from lobate-stage *M. leidyi*, reverse transcription was performed using Promega’s Go-Taq 2-Step system with oligo(dT) and randomized primers as per manufacturer’s instructions (Promega, Madison, WI).

### Poly(A)-tail length assays

cDNAs generated using the GB135 adapter as described above were used as templates for amplification and analysis of poly(A) tail length as per [97, 98] with slight modifications. Briefly, 2 μl of reverse transcription reaction were used directly as template for amplification of 3’ends using the GB136 T5 primer and a gene-specific primer (See Table 1 for utilized primers and annealing temperatures). Gene specific primers were designed to target sequence within ∼150 nucleotides from the furthest cleavage and polyadenylation site for each gene according to predicted gene models, EST reads, and RNAseq reads available for *N. vectensis* (https://mycocosm.jgi.doe.gov/Nemve1/Nemve1.home.html, [169]; https://genomes.stowers.org/starletseaanemone, [170] and *M. leidyi* predicted transcripts (https://research.nhgri.nih.gov/mnemiopsis/; [106, 171, 172])). 5 μl of product from 35 rounds of PCR were analyzed for each time-point in by 2% agarose gel electrophoresis (Top Vision, Thermo Scientific, Waltham, MA). Each result is representative of a minimum of two independent replicates.

For generation of 3’-end PCR products used in Amplicon-EZ sequencing, cDNA was synthesized as described above and 3 μl of reverse transcription reaction were used as template for 35 rounds of amplification using *Nv_cyclin1*, *Nv_cyclin3*, and *Nv_c-mos* forward primers with partial Illumina adapter sequence on their 5’-end, as well as primers including the GB-136T5 oligo sequence with a second Illumina adapter sequence, as per recommended by the sequencing service provider (Genewiz, South Plainfield, NJ; see Table 1). Amplicons were purified using DNA Clean & Concentrator-5 columns (Zymo Research, Irvine, CA), eluted in water, diluted to a concentration of 20 ng/µL, and shipped for sequencing.

### Computation analyses of poly(A)-tail sequencing Amplicon-EZ data

We used NGmerge version 0.3 [173] to merge overlapping paired-end sequences and correct erroneous and ambiguous base calls in our resulting poly(A)-tail sequencing data. We retained only those merged sequences that contained the 19-base-pair linker sequence. We masked the linker sequences and then applied a k-mer strategy (k=20) to mask UTR sequence present in the merged sequences. We considered poly(A) tails to be the bases between the masked UTR and linker sequences and calculated statistical data (*e.g.,* composition, mean and median lengths) from these predicted tails. All command lines and scripts used to identify tails and compute statistics are available in our GitHub repository (https://github.com/lrouhana/cpeb_evolution).

### Direct sequencing of N. vectensis mRNAs using Oxford Nanopore Technology

An mRNA library was created using Nanopore’s direct RNA sequencing kit (SQK-RNA002) from extracted total RNA of unfertilized *N. vectensis* eggs per the manufacturer’s recommended instructions (Oxford Nanopore Technologies, Oxford, United Kingdom). Sequencing was performed using the minION device. Base calling and quality control of minION’s .fast5 files was performed using default parameters on guppy. The resulting .fastq files were compiled into a reference .fasta file (Supplementary File S3), which was used to identify relevant reads via BLASTN, which were later aligned to the reference cDNAs in CLC workbench.

### Translational assessment of injected luciferase reporter mRNAs using N. vectensis

Firefly luciferase and NanoLuc luciferase mRNAs were generated from pT7-Luc and pT7-Nanoluc vectors templates [91] using the mMessage mMachine T7 Ultra transcription kit (Invitrogen, Waltham, MA). Templates were linearized with BglII and cleaned-up using DNA Clean & Concentrator-5 columns (Zymo Research, Irvine, CA). For polyadenylated mRNAs, recombinant yeast PAP from the mMessage mMachine T7 Ultra transcription kit was used on half of the *in vitro*-transcribed mRNAs, the other half was left without a poly(A) tail for comparison. After testing serial dilutions of mRNA concentrations (Supplementary Figure S3) it was decided to co-inject Nanoluc mRNAs (+/-p(A∼350)) with Luciferase mRNAs (p(A0)) as loading controls at a concentration of 25 ng/μl (each) into dejellied *N. vectensis* eggs and zygotes as per Layden et al. (2013). Nanoluc and Luciferase signals were measured in 96-well plates, from four groups of 5 eggs or embryos, 6 hrs. post-injection using a Synergy HTX Multi-mode Microplate Reader (Bio Tek Instruments, Winooski, VT) and the Nano-Glo Dual Luciferase Reporter System (Promega, Madison, WI) as per the manufacturer’s instructions. Luminescence from NanoLuc reporter was normalized to co-injected firefly luciferase signal and the average ratio of normalized NanoLuc signals were compared between groups of samples injected with polyadenylated and non polyadenylated Nanoluc mRNAs. Levels of reporter mRNAs in eggs were measured 6 hrs post-injection by RT-qPCR using the GoTaq 2-Step RT-qPCR system as per manufacturer’s instructions (Promega, Madison, WI) with random primers for reverse transcription and oligos listed in Table 1 for the qPCR step.

### Analysis of *MlCPEB1c* expression by *in situ* hybridization

Partial *MlCPEB1c* (ML05854a) sequence was amplified from *M. leidyi* cDNA using primers listed in Table 1. Amplicons were cloned into the pGEM-T vector (Promega, USA) and their identity confirmed using Sanger sequencing. Riboprobes labeled with digoxygenin (DIG) were generated using this construct as template for *in vitro* transcription (Megascript SP6 transcription kit, Invitrogen, Waltham, MA) and diluted into a 1ng/ μl stock working solution in hybridization buffer. *M. leidyi* cydippids with visible gonads and not fed one day prior were fixed as previously described and stored in methanol until their use for *in situ* hybridization as per Mitchell et al [174].

## ACKNOWLEDGMENTS

We gratefully acknowledge the Sunset Inlet Homeowners Association in Beverly Beach and Marker 8 Hotel and Marina in St. Augustine for facilitating animal collections on their properties. We thank current and former members of the Martindale laboratory, in particular Leslie Babonis, Radim Zidek, Brent Foster, Dorothy Mitchell, and Camille Enjolras, for technical advice and assistance during completion of this project. We would also like to thank Sandra Loesgen for generous access to lab equipment. Research reported in this publication was supported by The Eunice Kennedy Shriver National Institute of Child Health and Human Development of the National Institutes of Health under award number R15HD082754 to L.R., an Allen Distinguished Investigator Award and a Paul G. Allen Frontiers Group advised grant of the Paul G. Allen Family Foundation to J.F.R. and M.Q.M, as well as grants from the National Science Foundation (IOS-1755364) and the National Aeronautics and Space Administration (80NSSC18K1067) to M.Q.M.. A.E. was supported by a National Science Foundation Postdoctoral Research Fellowship in Biology (DBI-2010755). The funders had no role in study design, data collection and analysis, decision to publish, or preparation of the manuscript.

## Author Contributions

LR directed the overall framework of the project and performed molecular analyses and injection of mRNAs; AE contributed RNAseq reads from *Nematostella* and anatomical analyses of *Mnemiopsis*, FH performed *in situ* hybridization analysis in *Mnemiopsis*, JFR developed programs and performed bioinformatic and phylogenetic analyses; MQM provided organisms for molecular studies and performed injection of mRNAs; LR, AE, FH, MQM, and JFR contributed to preparation of the manuscript.

