## Supplemental data for "Cytoplasmic polyadenylation is an ancestral hallmark of early development in animals"

### SUPPLEMENTARY INFORMATION CAPTIONS

#### Supplementary Figures

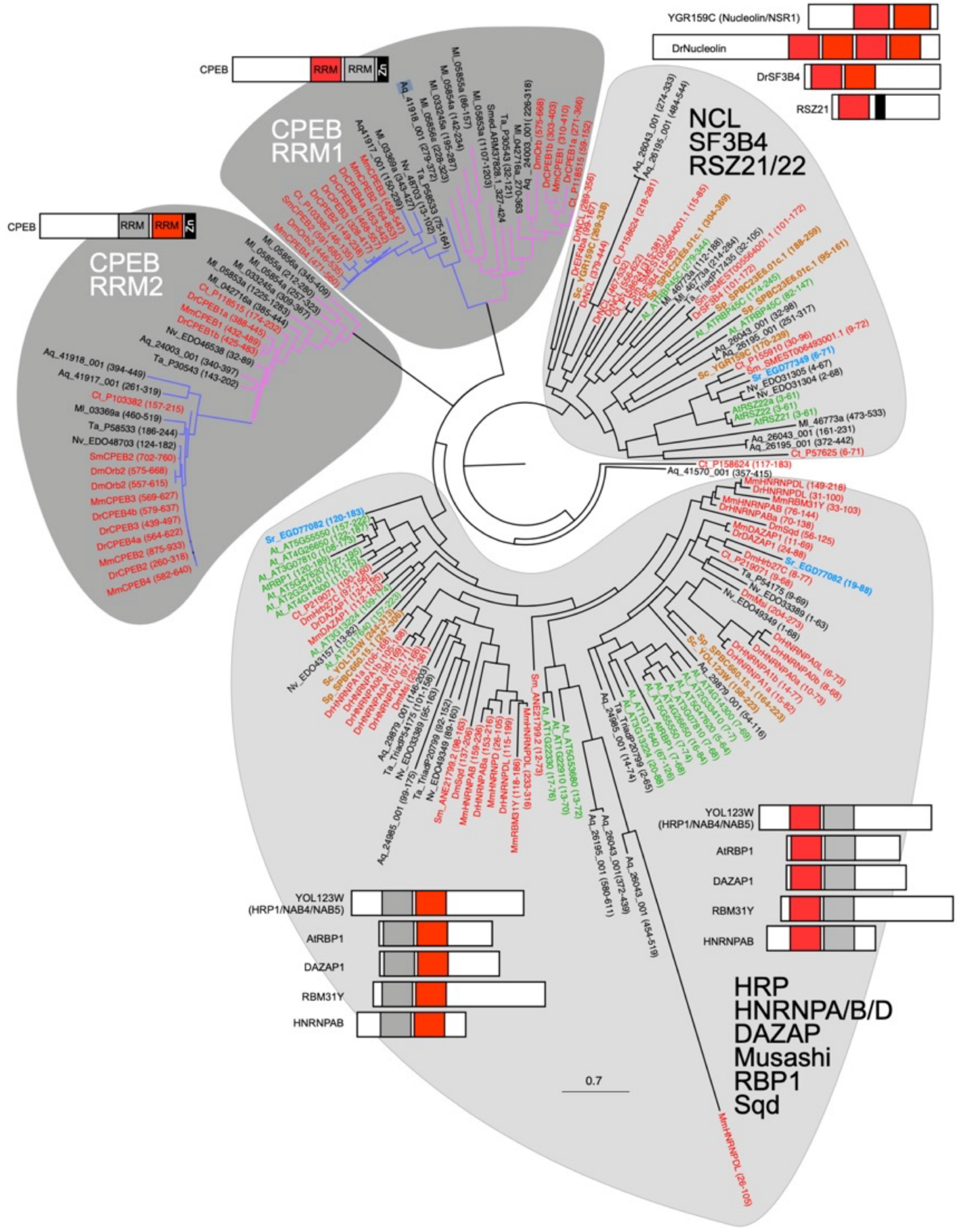

**Supplementary Figure S1**. Phylogenetic tree (as in Figure 1C) based on similarity to individual RRMs of CPEB1 depicts relationship between CPEB orthologs and close homologs.  Bar diagrams of representative proteins in each clade are depicted with the position of RRMs in the clade indicated in red boxes, other RRMs in gray, and ZnF in black.  Stems corresponding to CPEB1 (magenta) and CPEB2 (blue) sequences are highlighted.  Sequence identifiers are indicated with amino acid positions of RRMs in parentheses.

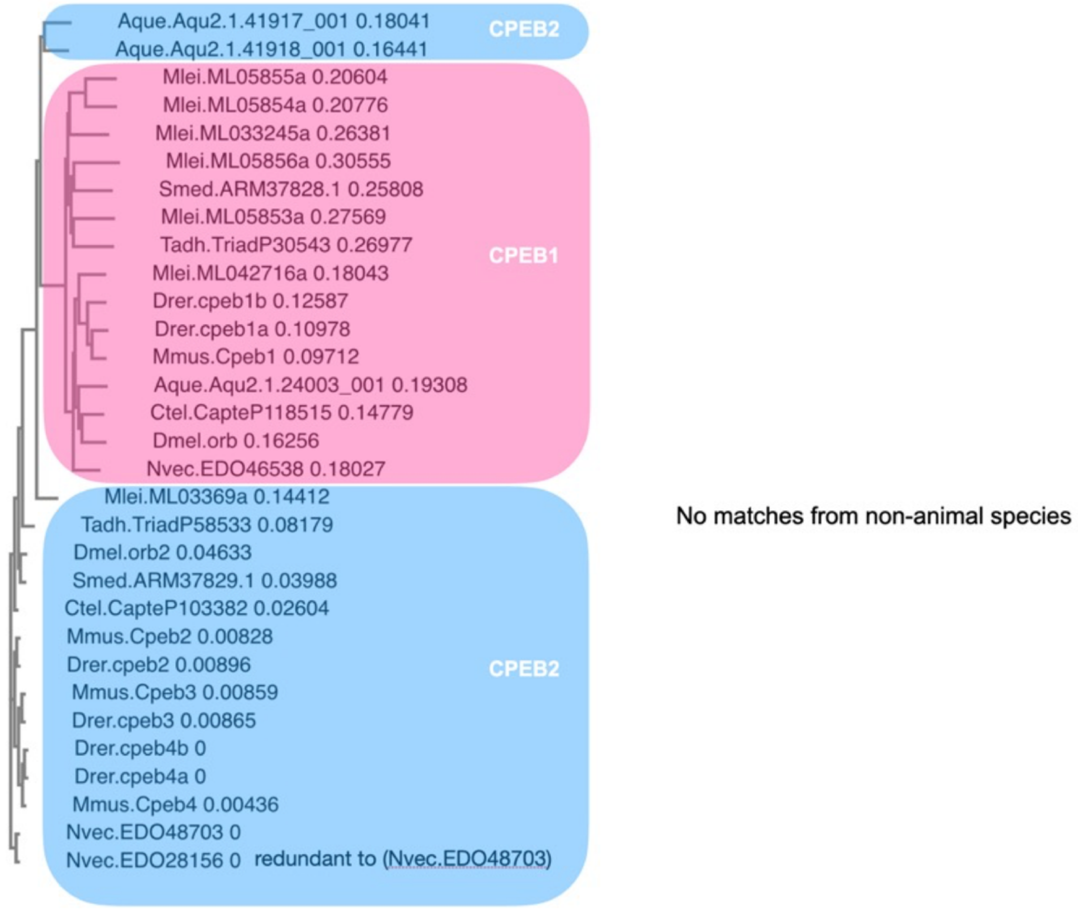

**Supplementary Figure S2**. Phylogram showing relationship between CPEBs from the five major animal lineages based on neighbor joining association of sequence matches to CPEB Zinc Finger motif BLAST searches. Sequences corresponding to CPEB1 orthologs are shaded in pink. and those corresponding to CPEB2 orthologs in blue. No hits were found in non-animal sequences.

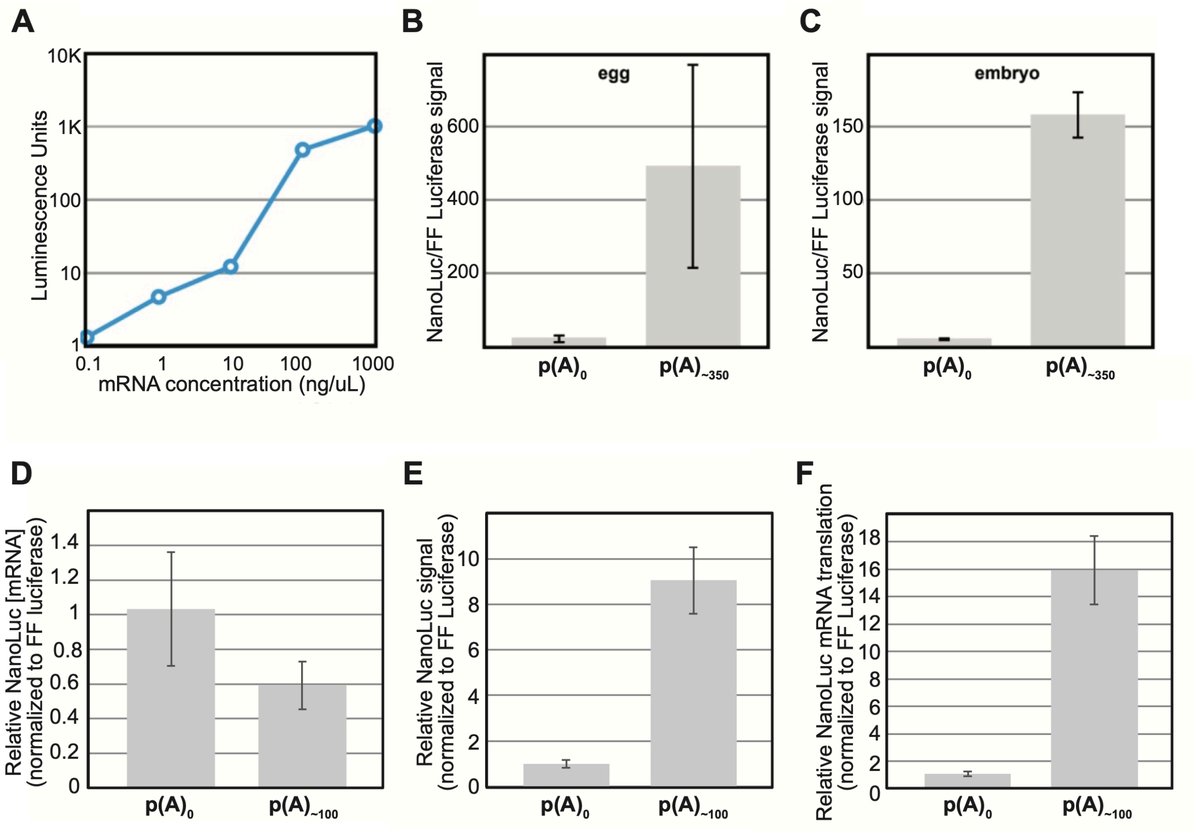

**Supplementary Figure S3.** The poly(A) tail enhances expression of reporter mRNA in *N. vectensis* eggs and embryos. **(A)** Luminescence units (*y*-axis) generated from eggs of *N. vectensis* 6 hours post-injection of different concentrations (*x*-axis) of *in vitro*- synthesized unpolyadenylated firefly luciferase mRNAs. **(B-C)** Translation level of unpolyadenylated (first column) and polyadenylated (second column) NanoLuc mRNA relative to unpolyadenylated firefly luciferase mRNAs 6 hours after co-injection [25 ng/µl ea.] into *N. vectensis* eggs (B) and embryos (C). **(D-F)** Results from a single batch of injected *N. vectensis* eggs reveals close relative abundance of non-polyadenylated (poly(A)$_0$) and polyadenylated (poly(A)$_{\sim 100}$) NanoLuc reporter mRNA 6 hours post-injection (D), whereas NanoLuc luminescence activity in the same batch of injected eggs was 16-fold higher for polyadenylated transcripts than poly(A)$_0$ counterparts (E and F). **(D)** The abundance of injected NanoLuc reporter mRNAs is shown normalized to co-injected Firefly Luciferase (FF Luciferase) mRNA reporter without a poly(A) tail. **(E)** NanoLuc luminescence activity in eggs relative to that of FF Luciferase in eggs from the batch as in (D) 6 post-injection of reporter mRNAs. **(F)** Relative levels of poly(A)- and poly(A)+ NanoLuc mRNA translation calculated from the ration of normalized protein activity in (E) to corresponding normalized mRNA abundance in (D).

>XM_001629488.2 (Nv_c-mos)
GTGGGTATGTCTTGCTCCCGGGACCAAGTTTGCAGTTGGACTCAAAATTTGCAAGACATTGACCGACTCGGTCTTCG
TACAAGAAGCAATTCCAATGAGTCCAAAGAAAGATAAGAATGATAAAAGAAACCGAAGTAGCATTTACGAGTAGTTT
AGACGTCAACGAAGTGACGATTCTCAAGGTTCTAGGTTCCGGCGGGTTTGGAACCGTTCACGAAGGCATTTACAGAA
ATCAGAGAATTGCACTGAAACGATTACACACTAACACGAAGAACAAGAAAGCAGCCATGCAGAGTTTCCAAGCAGAG
TGCAGAAACGAAGTGCTGTCACTAGACCACCCAAACATTATCAAGACGTTAGCAGTTTCAAGCGCGTTTACTCTAGA
AGACGAACCTTGCCTTCTAATGGAATTCGTCAGCACTAGAACTCTTCAGCACGTCATCGACGACACAAATGAGAAAA
TGGACTTTGCAAGGAAGTTACGCGTCAACAGTGAAATTAGCAACGCTCTGGTATACATCCACAACAGCGGGATTGTA
CATTTGGACTTGAAACCTGCAAATGTTCTGATAGCTGATGACGGGAGCTGTAAAATAGGAGACTTTGGATGCTGCCA
GTTCGTGGATGACCAGCCCAACACCCCTACGAGGTCTTACCTCACAGGGACCTTCGCTTACCGGGCACCGGAGCTTT
TGCGAGGCGAATCCCCGACCTTTCAGGCCGATGTCTACTCGTTCGCTATCTGCATGTGGCAGATCTGGACGCGTGAA
GTCCCGTACAAGTTGCAGAATCACCAAGTCGTGATTTTCCGTGTGGTCGCTTGTAGCCTTCGACCGCAGATTCCCAC
CGGTAACGAGATAGACAATCGGTACAAGGAGTTAATGACGAGCGCTTGGGCGGGAAAACCAACAGATCGGCCAACCA
TGGGAGACATCAAGGAAACCTTTGCAAACTGGAGAATCTGAGAAATCTTCGAGCATTTATTTTCTAGCTGGGCTAGA
ACTTAACCATAGATTAGTGTATTAGAGTATTTATTATATAACAGGATGTTGGAGCAGGTCTCTATAGATATAGTACT
AATTATAGAATTAATTATTATGCGTTCAACAAAAAAGCAGATAAAGTATTTTAATACGAGTGATTACAACAGAAAAA
AAACTTTTATTGGAAAGAAGCAGGAATTTTGAAAAAGAATTATCTGGAGGCTTTGTAATAAGAGTTTGTTGGAACAG
CTCGCTAAGAGTTCAATTATCTTGTGTAATTTATTCCAGAGTGAAATGACCTAGAGTGAAATTATTTATTATATAGA
GAAATTATCTATTTATTTATTTAGCAACATATATCATTAAGTGACAGAGATATATTAAATGCATAAAGCTTCATTAT
TTCAAAGTGAATTATTTATTATCGATTAATTTGATCGTTTATTCATTGACTGATTTCTGTATTAAATTGCTTATATA
TTTATAGGTATATTTATCTGTTATTCAAATAATTATTTGTTTTCTTATCCATTTATTTGTATTTTTTCTTATTTCAA
GTCAGTCTACACAATATTTAATTGATTAATTAATTAATTTCTGTGCATCTATTCATTTATTTATTTAGACTATCTGT
TTCTTCGTTCTTTCGTTTATTAACTTACTAACATACTCATTTATTCATTTATATATTTGTAAGCACTTCATTTATCG
ATTTGGCCATTTGCGCGTCATAGTCATCGCAATTAATTAAAGTCTTCCAACCAACATAACAAACTAAACATTTCTTG
TAACGGAAGTAAAATTCAATGAAACTGAATATAATGTAGTGTATTCCTATTTGCTGTGCTACTTTATTTTGGCGATG
TTATGAAAATAAACTTCTGTATGGTGCGTAA

-3’UTR region in amplicon: 178 nts.

>XM_032362260.1 (Nv_GLD2)
AAACCTATTGCGCATGCGCACATTCGTTCAATAATCAACATGTCGGCCGAACTCTAATGGAAAGCTGTGTACAGTAG
CTCTAAAAGTTCTGTACGAAGCAGAGAAACTCAACCATTGACAGAGCTATAAGTGTGTTTTGCTGGTAGTCTGATGC
TTGCCATGGCTCACCATAATCATTATTCTGAGAAAATTAACCTTAAAAGGCAACATTCACTTGTGTTTTGTGATACA
ATTGAGCAGCAGATGCACGAGGATATAACTCGAAAAAGGCAAGCTACAGATTCAGGTGTTTTCAAACCCAATCAAGC
AAGAAATTACCATTCTTATCATGATTTCAATAAACCTGAACAGATGCATAGCCACGCTTTTGATACATGTACAGGAC
CTAATAAACAGAGAATTGGAAGTTTGCCAGACAATTTCTCAAACTCACCAAGCTCTAATAGAAACTCACTGTATATT
AGAAATACCACAAATTCCAACAACATTGACAATTTAACAAGACAAATGGGTTCTTCATCAGAAAGCAATAATTTGGT
TGCTCAAAATAGGTCTCCATTTCCCCATCCTGTCAAAAAAGCTCCTAATTTTTCTGTCATTGAAAAAATAGTCCCAG
GACACAGGCAATTCTTAGATAGACCTTTTGCTCAAGCCAGAAGACAACTTGCAGATAAAAAGCGGCAAAGTTACCCA
GCACAAAATGAGCATGGACACACAATTAGGAACGATAGGCGTGCATCGTTGGGGTCTCCCCCAAGATTCAATGACAA
GGGGCCATTAAAGAGGCGGCATTTCTCGGAAGTTACACCCTCCAGCTATGATGGCTACAGTAACAAAAGGTTGCGGC
ATGGTAGTGGACCTCCAGTGGGGAATGCTCCGTGGTCTGATGAGCAACAATATTCTCCCCATTGTAGTTATGAGACT
CCTTTTAACATTGATACCTTGAGCAATGAGATTATGGCTGAATCTGAGAAGAATCTACAAACAGAAGCCACTCTGGA
AAAGAAAATGAAACTCAAGCAAGCACTTGAGAATGTTTTCCAAAGAAGCTTCCCTGGGTGCTCTTTACATCTGAGTG
GATCATCTGTGAATGGACTAGGTACAGATGAAAGTGATGCAGACTTTTGTCTCATGCTCACACAATGGGGTGAGATT
GACCAAAAACAGGAAGCAAGAAGAATCCTGATGATGTTGAATCGCATTCTTCAATCTTGTGATTTTATAAGAGAAAA
CCAAGTGATCTTTGCCACAGTACCTATTGTAAAGTTCATTGATGCTGTGAGCAATTGCGAGTGTGATATCAACATAA
ACAACCATGTTGGAATCAGGAACACGCATCTACTGCGCGCCTACTGCCTGGTTGATAGTAGGGTTAAGCCTTTGATT
ATGATCGTGAAGAAATGGGCGAAGAAACATCAGATAAACGACGCTAAGGATGGAACACTCTCCAGCTACGCCCTCAG
TCTAATGGTCATCAACTACCTGCAGTGTGGATGTGCACCTCCTGTTCTTCCATCTTTGCAAAAAAAGCACCAGGATT
TATTCTCACAGCATCGTGATGTCACCAAACTGTTGGAGGAGGACACTGCAAGTGTTTTGCAGAAACGAAGGAGTTTC
ATGCCCAGGAATAACCAGAGTGTTGGGTCCCTTCTTGTTGGTTTCTTCCAGTACTATGCCAACACTGTCAACTGGGA
CAAAGAAGTGCTGAGTGTGCGAGAGGGGGCACGTTGTCCACGAGATTACAAGTGGAGAACCAAGATGATGTGTATCA
ATGAGCCATTTGATGGGAATAACGTTGCTAAGACTGTCTACATCAAAGCCAAGTTTAAAACTATCAAAATGAAGATA
AGCCTGGCAGCCCACACTTTGCTGGTCTCTCCTTCACTGAAGAGCATACTGTAAATACTGCACAAATCAGGAAACCC
AGTGTCTTTAGTCACAGCTTCTAATGCTAAACAAAAAGAAACCAGATGACAAAAGAGCTTGAAATAGGGGAAGAAGG
GAGCAGAGGGCTGGGAGGGTATGGAATGAGAGATGTCTGATCACAAATATGCTTGGCTTGGCATGATTTTCATGCCT
ACTAACAGCAATGCTCAGATTTCAGGAAATGGTGAAAATATAAGAACAGTTGGAAGGGGACAGGATGGGGAGTTGTG
GAAGGAGGTTGAGAGACAGCATGATCAGCAGAGTAAGGCGGAGGCAGAGGGTGCAGCATGATGGGGAAGGGGATAGT

AAAGGGGGTGGGGGTGAGAGTAAGGCAAATGAAAGTCAACAAAAAGAGGGTTCAAACAGAGGTTAAGGGTTGATAAG
GGAGGGGATATGAAGAGGTTTGGGAGGCTAAAGGGCATGTGTAAGAGGAAGAGGGACAGGAAGGAGGTGGAAAAGGG
GCAAGTTGTTATTTGAATGGGTACAGGTAGGATGCGAGGTGTGAGGGGAAGGGGGTTGGGAAGGAGCTAATGTGTGA
GGAGAAGGGGTATGGAAAAGGGCGGGAAAGGGGCAAGGTGTTATTGGAAGGGGTAGGGCGCGAGGTGTGAGGGGAAG
GCAGACGGGAAGGAGTAAAGGTGTGAGGGGAAGGGGGCATAAGGGAGTCTGAAACAGGCACTGTTAGAAGGGTAAGG
GAATGGCCAAGTGTGTAGTGTAGAATGTAACAGGACAGAGGGTACTAAAGGGAATTTGAGGATATGAGGAATAGGAT
GAAATGCTGAAGATGACATGCGCAGAGAGTGGTAACAGGAGAAGAGGGGTGTGGCAGTTAGTGAGCTGAGGTTAGTT
AGTGAGAATGAGGTTAGTTAGTGAGGGTGAGGTTAGTGAGGATGAGGTTAGTTAGTGAGGGTGAGGTTAGTGAGGAT
GAGGTTAGTTAGTGAGGGTGAGGTTAGTGAGGTGAGGTTAGTTAGTGAGGATGAGGTTAGTTGGAAACGTGTGATAG
CTGTCAATTATGGGTGAAGTGAATATATAAAGGAAGACTTAAATGCCAAATGCATAATGCCAATACCTCTTTACAAA
TGTTAAAAACTGTTATCGTTAATGGCGTTGATGTTCAATGGCAAGGCACAGCGTCTACTCCAGACACGAGAATCTGA
AAAAATGTATAGTAGGTATAATAAATGTATAGAGAGAGAATGTTTGTAATGATATAGAATTCAATAATTAGCTTAAC
TATAGATCATCATATATTTATTTATATAAAGGAGTTTATTAATAAAGAATATAAATTTTTAAAA

3’UTR region in amplicon: 170 nt

>NvERTx.4.87874 (Nv_cyclin1)
AGAAAATTCGCGCCTTCTGATTGGCTGATTATAAAGCAAGCGGGCTTCTTGAAATATCAAGAGCTGCAAACGTTCAC
TTCGTTCACCACATCGCGACTAGATTCGAACAGTGCTGTCGTATCTAGCTAAAGAACTTGCCAAAAAACTTCACGAA
CGTCAAAGAAAACTTCAAGGTATACACTTGAATACCTAGCTACTCTAAAGTGAAAGCTTAGAAGTAGATTCTTGTTT
CTAAGACACATTTTATAGTCACGAAATTGAATACACGAATTTAGGATTTTACCAATTTGATTCAAATACTTTTTTAA
ACATGTCGCTGGCATTTGGAGGCGCTTACCAGTTTTCAAATAACTCGAACCAAGAGATTGAGGAAAATAATACCTTT
AAAAAAGCTAGGCTTGAAGATGGACAAGTGAAACTCACGAAGCGGGCGGCTTTGGGAACCATCACCAACAACGCGGG
AGTGAGAGTGCAGCCACACCGCGCTGCCAAGGTTGCCGTCTCTGGAAATGGAGGCCCAGTATTGAAGAATGACGAAA
ACGCGTTTAGCCGCGCTGGCAAATCCGCCTTCTCCATTCCCAACACTGCTCAACAGAGCTTCGCTATTCACGTTGAC
CAACAGCCAAACATCTCGAATTCACAAAAAGCTACAAGTCTAACTCAACAAGAGCCGGCCTTGAATCCTGCTGTAAC
TTCATTATCACGACCCGCACTGACGAATGTCTTCGTTGCGAGCGATGAAGGAAGCCTTGAATCTCCTATGGTCATTG
ACTCGGATTTTAGTAGCGACGATGAAGACATTGCAAGCAGCAGCCCGAGCAGAGACCAAATTCACAACATCGACTCG
GTAGCCGCCGACCCCATCCTCGGTGTACCCGAGTACGCGTCCGACATCTTCAAGTACCTCAAACAAGCAGAGCTCAA
TAACAGGGCCAAGCCAGGCTACATGCGGAAACAACCCGATATCAACAACAGCATGCGAGCTATCCTTGTCGACTGGC
TGGTGGAGGTAGCTGAAGAATATAAACTCCTTCCCCAGACCCTATACCTCACAGTAAACTACATAGACAGATTCCTG
TCTGCTATGTCGGTGCTGAGAGGCAAGCTACAACTTGTTGGTACAGCTTGCATGCTTCTCGCCTCCAAGTTTGAAGA
GATTTACCCGCCAGAGGTCTCCGAGTTTGTATACATCACAGATGACACTTACACAGCCAAACAGGTTTTGAAAATGG
AACAACTGGTTCTAAAAGTTCTCACCTTTGATCTGTCTGTACCAACCATTCTCAACTTCCTGGAGAGATTTATCAAA
GCAACCAATGTACCAGAATCTATGGCTCCCAAAGTAGAAGCTCTTGCAAGGTATCTCTGCGAGATCTCACTACTGGA
CAGTGAGCCCTTCCTGAAATATCTTCCTTCGACAATTGCTGCATCAGCGATCGTATTATCTCTTCATACATTAGGAC
TATCATACTGGAACAATACTCTGAGCCACTACACTGGATTTGAGCTTCATGACCTCCAAACATGCATACAAGATCTG
CACCGTAGCTTTGCCTATGCACCCAACCACCCCCAGCAGGCAACAAGAGAGAAGTACAGGAGTGCAAAGTTCCACAG
TGTGTCTAATCTCTCCCCACCTGACTGCTTGCCATTGGCATGAGAATCAAATTGCGTATAGATTTTTTACAATCCTA
ATACCATTTTATAGTATAGATAGAAACATTTTATCGTAATATTAATCAAGTGTAAAACAAGTCTTTTATTGTGTTGA
GTTTTTTACCCTAGCTTTTGCTTGTTTTTAAATGTATTGTAAACAATTATATTCACAATATGATGTACATATTATAT
GGTTTAGGGAACTTTTAATCTCTCCAAAAGAAACTGCATTTGAAATGCAAGTATGGCAGAGAAATCTTTTGAAAAAT
GTTTTGATAAATAATTCCCTATTCAGTGATTTCATACTATCTTGCATAACTTGTTTTATTTTCTAAATGACAGTCAT
ATAGTTTTAAATTATTTTTTAGTGAATTCTCAATGATACTGTTTAAACATGTCACCTATTGCCTATGCTTTATGATA
ACAGATGACAAAGTTATGCTGTTTTTTTTTTTACCATGTAGATTCCATATCCCAAGAGGTATTGTTCACCCATTGTA
ATATAATCCTGGATTGTACAGTATTTATAAATAAATGTGTAAATAATAACTTGGTTTATTTTCTTAACATGTTAAAG
GGTATCAGCTAGAGTGAGTTTTCAAAATCCTGAGTAGCAAAATGTAATTGTTTAAGTTTAATGGTCAAGCGATAACA
AATAGCACAAAGACATTACAG

3’UTR region in amplicon: 93 nts.
Predicted based on reference: 219 nts.

>NvERTx.4.58222 (Nv_cyclin2)
TCTACACTTCAAACGAAGAACTGATCAACAGCGAAGCAACACTGCAAGAGAACACCGAGATTTTGTCAAATTACACT
GAATTACCACAGAAATTTCTACATCTCTCTTGAATATACTGAAATGGCTGCTGTACGACGTCTTACCGCACAAACCG
TCCCAGCTCAAGAAAATGTCGACGTCCTCACAAAAGCTAAGCATGGCCAAAATACTCGCTTTGGACGCGCCGCTTTG
GGAGACATCGCAAACAAAGACAAGGCTGTTCTTCCCGGCAAAAGGATAGCTTTAGGAACGCGTGGTCTTACGAGAAA
TGAAGCTTTCACGGCGCTTCCCAAGCCCGAACGGCCGGCCTCAAAGCCCGAGCCCATGGACATGGCTGATTTTAGTG
AGGCGCTCAACGAATGCTTTCCTACCGATGTGGAAGATATCGATAGCGGAGATTACGATAAACCCCAGCTATGCGCC

GAGTACGCAAAAGAGATCATGCGATTTCTCCGAGCTATGGAAGAACACTACAGCGTTTCTCCCACTTACATGAATAA
CCAACAAGAAGTAAATGAAAAGATGCGTGCAATCTTGCTTGACTGGCTTGTGCAGGTCCACCTCAAATTCCGCCTTC
TGCAAGAAACCCTCTACATCACCATGTCAATTATAGATAGGTTTCTTGCTGTCCATCAAGTTTCTAAGAGAGAGCTT
CAGCTTGTAGGAGTTGGTGCTATGCTTCTTGCTTCCAAATATGAAGAGATGTTTGCCCCTGAGATAGGCGACTTTGT
CTACATCACAGACCATGCATACACCAAAAAACAGATCAGACAAATGGAATCCTTGATCTTCCGTAAGCTGGACTTCA
GTCTTGGCAAGCCACTCTGCCTTCATTTCCTCAGAAGGAACTCAAAGGCTGGAGCTGTTGGTGCTGAAGAACACACC
ATGGCTAAATACCTTATGGAGCTAACACTGATAGACTACCAGTCCATCAAGTTCCTCCCCTCTGAGATTGCTGCAGC
ATCTCTCAGCCTGGCCATGCGAGTTATGGGGAAAGGCAGTGAATGGACACCCACCCTTGAACACTACAGTGGTTACT
CTGAGAAGAAACTTAGCACATGCATGCAGAGATTGGCGCAGCTGGTTCTAGGAGCCAGGGACAGCAAACAGAAGGCT
GTGTACAACAAATATGCCAGCAGCAAGTTCATGAAAATCAGCACAATGTCTTGTTTGTCTACCTCCACGATAACAAC
TCTTGCAGCCCAGGACCAATCTTAGTTTATAATAAGCATAGAAGCAATTTATAAAGGGTTATAGGGTGGGTAGGGTG
GTTTAGGGGCAGGACCCATTCAGTAGACTCCCGTTCAGGAGAGTGTAAATAATGCCTGTAAACAGTGTCAGGTTTGA
AGTTATAAACACTGTGATTTCCAAGGAAATACTTTTTTTTAGTTTACTAATAACATTTTGATGTAAAAGTAATTTTA
TTGGAAACCACAGCTTCTGATATCATTTCATGAAATATAAATATTGGGATGGCTACATGGACCTTGTGTCAGTTACT
TTTTAATCACTAGTCTTTGTAAATATAGTTATATATTTTTTTTTCAAAATAAACCATTGATACTATTATATAGTCT
TTTCACTATGTGT

3’UTR region in amplicon: 96 nts.
Predicted based on reference: 116 nts.

>NvERTx.4.144958 (Nv_cyclin3)
GTGTGATGCCTAACCCCGGCAAGATTGAATAGAACGTCTGATCGGGACGCTTCCAAACAAGGATCGCATTCGACGAG
AAGTACATCAAAGCAAAGTTTTATCCAAAATTTTCTTACCATCTCTTATTATTTGGAGATATACATAAATTTACCCA
TTACTAGGCATTACTTTTAATCACTATGGTGAAAACGGCTCGAAATAGGCAATTCTCTGGGATTTTTCAGCAAAAGA
AGAAGGGATCCAAGAGTAAAGAATTGAGCGAGAATGGAGTCCTTGGAATTGGTGGGCCCGTAGAGCAAGTCAAAAGG
CTGGCAGCCTCACCACAAGAAGGCGAAGCACCAACCAGAAAGCGTTCTGCCTTTGGCGATATCACAAATGCTTTTCG
CCAGCAGCAGGCGGGGAAATCAAAGAAGTCCTCAGCTCAAAAGAAGCCCGAAACAGGAACGGAGTCTCAAAATGGCG
TACAAAGAAGAAAGACTCGAAGCTCTGGTGACCTTCCTGACTTCGAACTTCTGCCTTCGTCGGGTGAAGTCACGAGC
TCTCAGGAGTCTTCAGAGTCGAGCATTGACGTCCTAAGCGACATTTGCAGCGGCGTAGAGTTGACTGACAGCCAAAA
AATAGACAGCAGTGTAGGGTCTGAAATAGATTCTTTATTGAAAGAGCTGGATGGGTCAATAAGTTCAGAACAAGTTG
AGAACTCCTCACATCTTCCACCTGATGTTGTTGACATTGATGCTGACAAGACTGACCCATTTCAAGTTGCAGAGTAT
GCAGAAGAGATTTTCTTAAACATGAAACGAAGAGAGAATCTGTTTCCATTGGAGCTGTACATGGAGACACAAAAAGA
ACTAACCATAAGCATGCGGGCTATCCTAGTTGACTGGCTGGTTGAGGTTCAGGAAAGCTTTGAGCTGTATCATGAGA
CCTTGTACCTTGGTGTGAGAGTTTTAGACAACTATTTAATGCGTAGCTATGTGGAGAGAGAGAACCTCCAGCTTGTG
GGAGCTGTCAGCCTTTACATCGCCTGCAAAGTTGAAGAGAGACACCCACCTTGTCTTGATGACTTCCTCTACATCTG
TGATGATGCTTACCAGCAAAAGGCCTTTGTTGCTATGGAGAAGAAAATTTTGAACAGCTTGGAGTTTAATATCAATA
TGCCAATACCTTACAGATTCCTTAGGCGGTTTGCAAAGGTTGCCTCTGCTGATGTTAAGACTTTGACGTTGTCTCGT
TTTATCCTTGAAACCACCCTCCACCATTACAAGTTCATTGTGCATAAACCATCCTTCCTAGCCGCTGCCTGCCTTAG
ATTAGCCCTCCGCATGAAGGGATGTGATGACTGGACACCAACTGTTGTTCATTACACTGGATATTCTGTGGCACAGC
TTGATGGGTGTGTTATTGAACTCAATGAGATGATCTCTGAGCCCCCCAAGCAGAACCTGATGACTGTCAGAAACAAG
TACTCTCACAAAGTCTTCCATGAGGTTGCTTTGATCCCACCTTTGGACTCCCTAAATCTCTAGATAATTATTTTTTT
TATTTTGGTCACCTTTTTATTAGCCTTTCTATTGGTTTAAGTAGAAAGTTTTACAAAGGATTTTAAATAACATAATG
AACTTTACATTTCAACCATCAGAAACATTTCCTGCAAGGCTACTATTTGTGGCATTTAGTACTCATTCAAGGGTATC
TTAATCAGCTTTAAAAAATAAGGTATCAGATTACCCAATTTTAGATTACCCATATTGAGAAACATATCATTTTCATC
AAGAGATAACATTTTATTTAAATTTATTTGATAATGAACAGAGCAGCTTTGATTAACAAGCAGGAATTGTATATAT
TGTAAATATTCAACACTATCAACAAGTAAAGTTAGCTGCATAAGGGGAC

Observed 3’UTR region in amplicon: 224 nt.
Predicted based on reference: 232 nt

>ML4553_cuf_38 transcript (ML455312a; M. leidyi cyclin homolog 4)
GCAGCCAAGTTCTCAGTGTTCAAATTGGAGCTGATATAACTATATAACATGCCCCTGGTCTACACCGAGAACAACGA
TGAGAATTACCCCCCTATAACCTCCATGTTACGGAAGACAAAGCGGGACGAGCCCCGTCTGGCGGGGAGCGCTAGCA
AACGTCCAGCTCTCGCTACCATCAGTAACATTCAGACTTCTAGCAATGATCTCGGAAAGCAAATCAGGCAGCGTCAG
ACCAACCCCTTAAGTATCAGAGTGGACACCAAGAAGCCGTCTCCCAAACAGAAGCAGATCGCGTCCCGATCGCAGCG
CACGTCCTGTACTAATTCGCGCGTCACCTCAGTAGCCGGTGCCCTGAGACAGTCCACTCTGGCCACCTTCACTACTG
TATCCTCCTCCAACCCTGCCTTCCCCATCTTTAAAGACCGAACTTCCACGAGCTCTACCACTTCCGCTCCTTCCTTC
GCTGCTCTCAGAGACAGAACCAACATCCCCTCTGCATCTACTCCATCCTCCAGCAAAACCTCAGAATGTCCGATGTC
TGTAGACACTTCATTGCGTGTTCCGCTTATCCCTCGCCGTCGACATCCTTCTTGTGCCTCCGCTCGCAACATGGAAA

TCGACGATTCTTCTAACGTCTCCGACATGAGCATCGATCTTATCGTCGAGGAGCCGAGGATCAAGAATATTGATGAA
GGAGCAAGTCTTGAGGAAAGTCCGGAGTACGCTGAGGATATCGTGAACTACTTGCGAGTTATGGAGGAAAAGTACAA
GCCTAAACCTGGGTACATGAACAAGCAACGTTACATTAACTCCGCCAACAGATCCACCGTCATTGACTGGATGTTCG
AGGTCTGTGACGAATTCCACCTCTCCCAAAAGACATTCCAGCTGGCTGTCTCCTACGTAGACCGGTTCCTCTCCAAG
ATGTCGATGCCAAGAAAAAACCTCCAACTTCTAGGAACTACCGCTCTCTTTATCGCCTGTAAGGTGGAGGAGATCGA
GGTTCCTGCCGGAAGTGCGCTGGCTGCTCGATTTGTCTGGATTACTGATAACACTTACACCGTCAAAGAGTTGTTCA
AGATGGAAGTTCTGATCCTGGAGAAACTTGGCTACGAGATCAACTCTCCCAGTACTCTTTCCTTCCAAGATCGGTTC
ATCAAAGCTTCCGGTGGTGACTGTACTGAGCAGTACTTCACTGAGTACCTGTGTAACCGATCTCTGATCGCCGGAGA
CAAGTTTCTGAATTACAAACCCTCTCACGTGACCGCTGCTGCTATCGGTCTGTCACGTGCTCTCCAGCGACCCTCCG
AACCCGTTTGGACCCCGACCCTGTCTCACTACACGAAATACACCTACCCCGAGATCAAGGAATGTATGTCCGATCTC
CTCCAACTCTACAAGTGTGACTTCAACTCTGAAACTCGGCGGTTTGGTGCTGTCTGGAACAAATACAACCTCCCTCG
ATACATGTTCGTGCTACAGAAGAAACCCATCGACTCAGTACCCGCACATCAATGAAGTTTCTCAACAAAAATCCCAT
CAAACAAGTTCCTTGTCAGTGTTCCAGTTCCCGTCCCACATTTGGCTGTCAATTAGCTAATTGTTTAATTACCGGCT
AATTGGCACGCCAGATGAAATAAGATTCTTTCAGAATCACCGATCCCAGGATCGAATCTTTATAAACTCTTGTGTAA
TTTTCAGTCGAACTCTCAGTGTTCCTAATATGTTTGTTTTGCTTCAGTTGAACGAATTTGATAATTTTGATGGAATT
ATTCAATTAATTTACATCAGCAATCAATTTATTACTCGAGGTTTATTTTTGTAGTGCAGTAGATAAACTGCCTGACT
AATAAGTTGACAGACCATATTTATTTGACTGCAGTCTTACATTTGTTGGTTATTTTACAGAATTTCATTTTTTTTCA
GCTCAACTTTCCGAACAAAACAAACTTTCTCGTGTGAAAATAGGATTAATATTTGACCAAGAACCTGACGAATATAG
TTACTTGAATGATGGATAGGACCTAGCTAATATAGAATCTAGCTTTAATTAGTCTCGAAAATATTTTCCAAATGACG
AAACAGTTCTGTACATATTTTATATGATCTGTAAAATTTTGGAGTCTAAACGAAGGCCCAAAAAGGATGAAATCTAA
CATTTTGTACCATTCTGTACGTATTTTATACGTATTTGTACGTATTTGAATTTTCCACTTCCCCTGAGAGCAACTTA
AGAACATTCTTTTTGGGCCGCATCACACTGTAGTCTTTCTCTTAACCTTTTGAGTTTTTATCGGTATCCCTACCTTA
TTAGTAGAATAGTGTTCGCGTATGTTGTATTTGTATATAATGATAATTTTAGTACAGCCCTTTACGGTTAGCTTGGG
CAAGTTGAAGTTTGAATGGTATCAGTAACAAGTATGATATGATAATTAATTGATTATTAATTATCCAATTGGTTGAT
ATTACTTTTGTTACTTGACTCAACCCTC

3’UTR region in amplicon: 191 nt

**Supplementary Figure S4.** Reference cDNA sequences for targets of cytoplasmic polyadenylation identified in this study. Sequences obtained from NCBI (https://www.ncbi.nlm.nih.gov/; XM_ prefix), SIMRbase (https://simrbase.stowers.org/; NvERTX prefix), and the Mnemiopsis Genome Portal (https://research.nhgri.nih.gov/mnemiopsis/; ML prefix) are shown. The predicted open reading frames (highlighted in gray), putative Cytoplasmic Polyadenylation Elements (UUUUA; red font), as well as Cleavage and Polyadenylation Signals (underlined) are indicated. Highlighted in yellow are regions validated from Sanger sequencing of cloned poly(A) tail length assay amplicons, along with the size of highlighted UTR regions in the amplicons. The predicted 3’-end lengths of reference sequences including regions not covered in Sanger-sequenced PAT amplicons (potentially due to alternative/rare cleavage and polyadenylation sites) are also indicated when present.

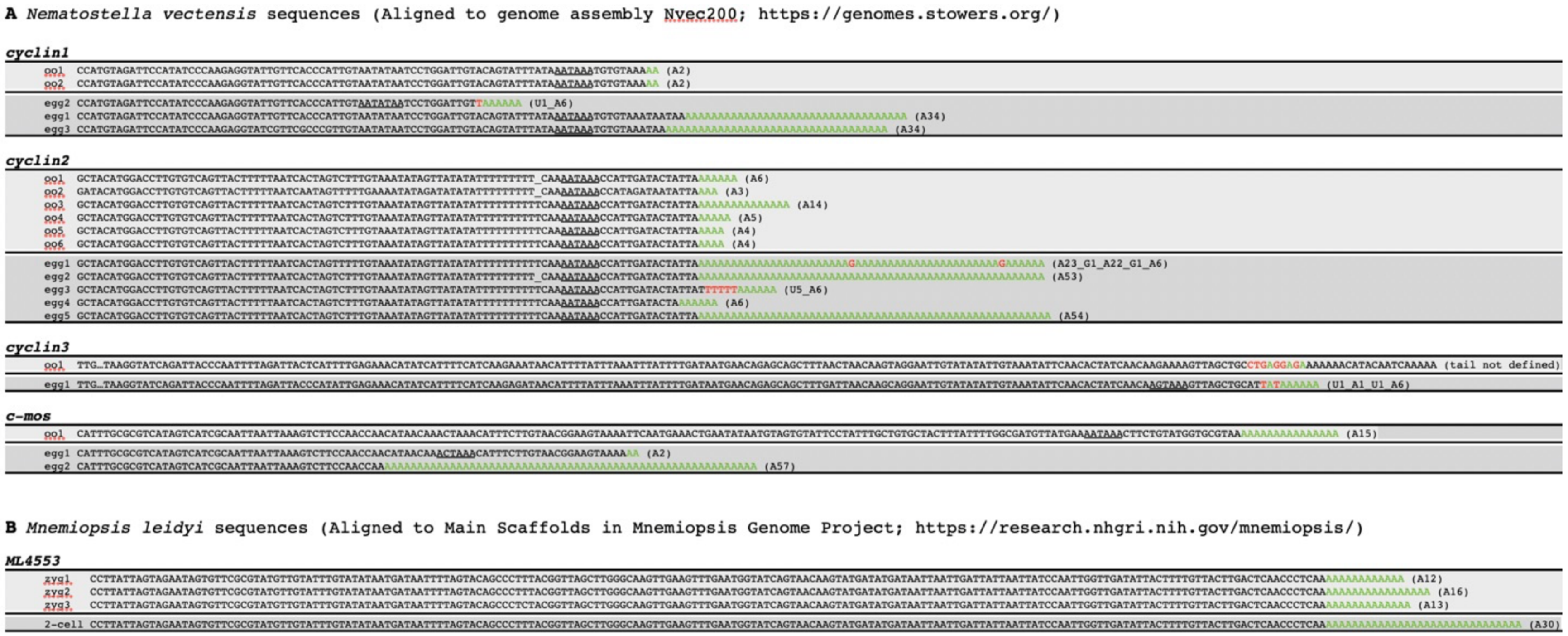
A Nematostella vectensis sequences (Aligned to genome assembly Nvec200; https://genomes.stowers.org/)
cyclin1
oo1 (A2)
oo2 (A2)
egg2 (U1_A6)
egg1 (A34)
egg3 (A34)
cyclin2
oo1 (A6)
oo2 (A3)
oo3 (A14)
oo4 (A5)
oo5 (A4)
oo6 (A4)
egg1 (A23_G1_A22_G1_A6)
egg2 (A53)
egg3 (U5_A6)
egg4 (A6)
egg5 (A54)
cyclin3
oo1 (tail not defined)
egg1 (U1_A1_U1_A6)
c-mos
oo1 (A15)
egg1 (A2)
egg2 (A57)
B Mnemiopsis leidyi sequences (Aligned to Main Scaffolds in Mnemiopsis Genome Project; https://research.nhgri.nih.gov/mnemiopsis/)
ML4553
zyg1 (A12)
zyg2 (A16)
zyg3 (A13)
2-cell (A30)

**Supplementary Figure S5. (A-B)** Sequence of 3'end amplicon cDNA clones from oocytes and eggs of *N. vectensis* (A), as well as *M. leidyi* zygotes and 2-cell stage embryos (B). Sequence that matches genic reference is shown in black font, whereas the rest is presumed to represent poly(A) tails (green font). Non-A nucleotides in poly(A) tails are shown in red and composition of tails summarized in parentheses. There is sequence downstream of the highlighted region at the end of the oocyte *cyclin3* read that matches genic reference. This is presumed to be amplification from a longer 3'UTR with indicated sequence (red and green font) that does not match our genic reference.

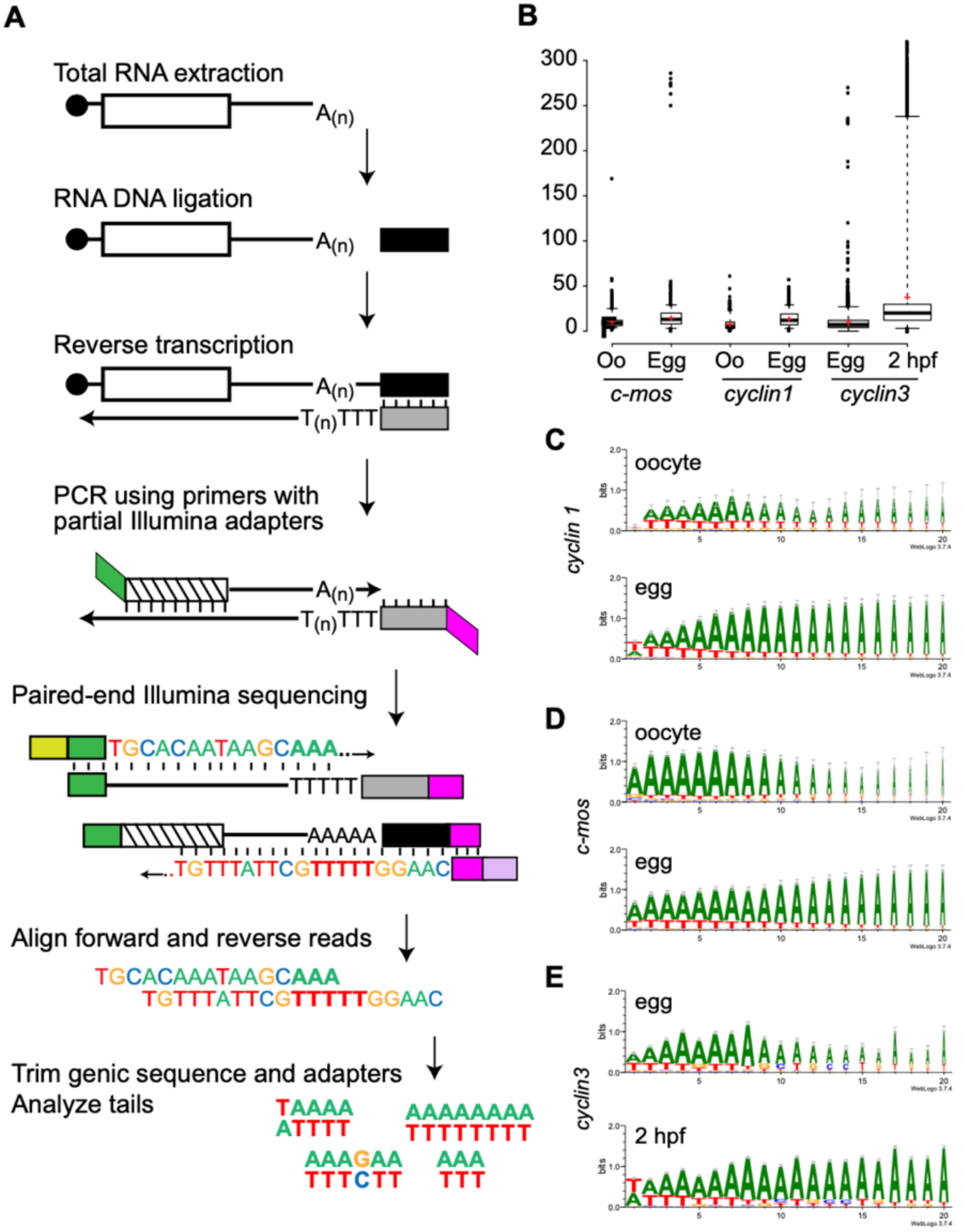

**Supplementary Figure S6. Poly(A) tail length and composition assessed by Amplicon-EZ sequencing. (A)** Generation of Amplicon-EZ sequence reads corresponding to 3'-ends of mRNAs involves the same initial steps as in (Figure 2A), but with modified primers that include adapters for Illumina sequencing (green and magenta) and is followed by alignment of paired-end reads and trimming of adapters and genic sequence. **(B)** Box plot of poly(A) tail lengths measured using Amplicon-EZ from ovary (Oo), egg, and 2 hours post-fertilization RNA samples. Thick horizontal lines represent medians; red crosses represent sample means; box limits indicate the 25th and 75th percentiles; whiskers extend to 5th and 95th percentiles; outliers are represented by dots; and width of the boxes is proportional to the square root of the sample size (n= 3249, 3285, 631, 2381, 12804, and 17482 from left to right of the plot). Plot was generated in BoxPlorTR (http://shiny.chemgrid.org/boxplotr/). **(C-E)** Sequence logos for the first 20 nucleotides of poly(A) tail sequence as obtained from Amplicon-EZ analysis of *N. vectensis cyclin1* (C), *c-mos* (D), and *cyclin3* (E) mRNAs. Sequence logos generated by the WebLogo 3 server (Crooks et al. [101]) for all sequences with poly(A) tails longer than 5 nucleotides.

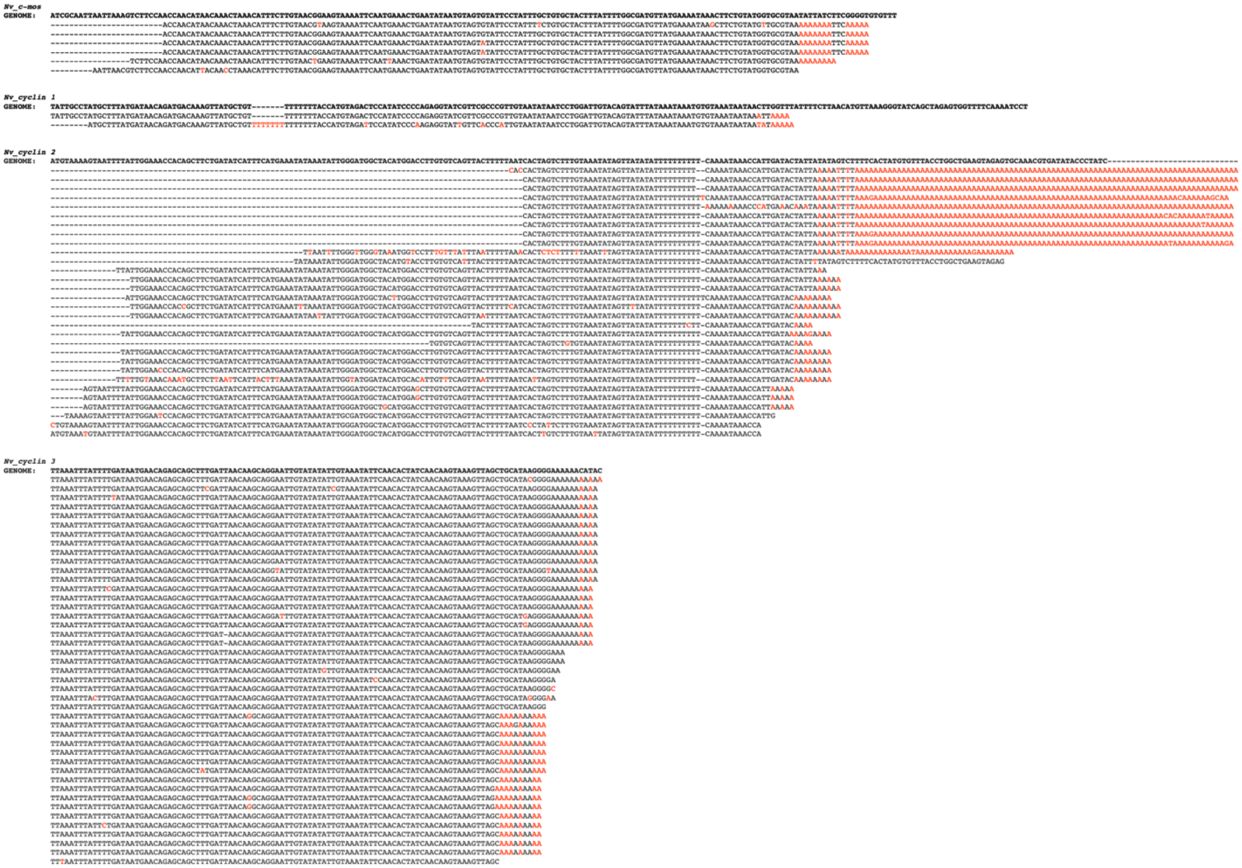

**Supplementary Figure S7**. Sequence of *N. vectensis* ribodepleted Illumina RNAseq reads (NCBI BioProject: PRJNA893363) aligned to corresponding genomic reference sequence of cytoplasmic polyadenylation targets (bold; Nvec200 Genome Assembly, https://genomes.stowers.org/files/pub/nematostella/Nvec/genomes/Nvec200/Nvec200.fasta). Positions that do not match the genomic reference are marked in red font.

| cyclin1 sequences | Length |
|---|---|
| UUAAAAAGAAAAAAAUAAAAU | 21 |
| AAGAAAAAGAAAAAAAC | 17 |
| AAAAAAAAAGAAAAA | 15 |
| UUUAAAAAAAUAAAA | 15 |
| AAAUUUAAAAAAAU | 14 |
| AAAAAGAAAAAU | 12 |
| AUAAAAAGGAAU | 12 |
| GAAAAAAGAAA | 11 |
| AUUUUAAAAA | 10 |
| AGAAAAAAU | 9 |
| AUGAGAAAU | 9 |
| AAAAAAAAU | 9 |
| GGAAAAAU | 8 |
| UUGAAAAU | 8 |
| AAAAAAC | 7 |
| AAUAGAA | 7 |
| AAAAAAA | 7 |
| ACAAAAA | 7 |
| AAAAAU | 6 |
| AAAAAU | 6 |
| AAGAA | 5 |
| AACU | 4 |
| AAAU | 4 |
| AAA | 3 |
| AAA | 3 |
| AAA | 3 |
| AU | 2 |
| AU | 2 |
| AU | 2 |
| AU | 2 |
| A | 1 |
| U | 1 |
| average: | 7.5625 |
| standard deviation: | 5.066891266 |

| cyclin3 sequences | Length |
|---|---|
| UAAUCAAAAAUAAAAAUAAUAUUAAAAAAUAAAU | 34 |
| AUAAAAAAAAAUAAAAAAAAAU | 22 |
| UAUACAAUCUAAAAUAGAAAAA | 22 |
| AAUAAAAAAAAAAAAAAA | 18 |
| AAUAAUAAUUAAAAAU | 16 |
| AAUUAAAAAUAAAAAU | 16 |
| AAAAAAUAAAAAAAU | 15 |
| AUUUAAAAAAAAAU | 14 |
| UUUUAAAAAAAAA | 13 |
| AAAACAUAUAAU | 12 |
| AAACUUUAAAAU | 12 |
| UACUUAGAAAAU | 12 |
| UUUAUUAAAAAU | 12 |
| UUCUAAAAAAAU | 12 |
| UAAAAUAAAAAU | 12 |
| AAAAAAAUAAU | 11 |
| AAAAAAAUAAU | 11 |
| AAGAAAAAAAU | 11 |
| AAAAAUAAAAU | 11 |
| UUAAAAAAAC | 10 |
| ACAAAAAAUU | 10 |
| AAAAAAAAAU | 10 |
| UUAAAAAAU | 9 |
| UUAAAAAU | 8 |
| AAAAAAA | 7 |
| AAAAAAA | 7 |
| AAAAAAC | 7 |
| AAAAAAU | 7 |
| AAAAAAU | 7 |
| AAAAAAU | 7 |
| AAAAAC | 6 |
| AAAAAU | 6 |
| AAAAAU | 6 |
| AAAAAC | 6 |
| AAAAAA | 6 |
| AAAAA | 5 |
| AAAAA | 5 |
| U | 1 |
| average: | 10.94736842 |
| standard deviation: | 5.959082863 |

| cmos sequences | Length |
|---|---|
| AAAUUUAAAAAAAUAAAAAUAAAC | 24 |
| AAUAUUAAUAAUUAAAAAAAU | 21 |
| AAUUAAAAUAAAAAUAAAAU | 20 |
| UAUUAAAAUAAUUAAAAAU | 19 |
| UUAUAAAAAUUUUCAAAAU | 19 |
| UAUUAAAAAAACAAAAAAU | 19 |
| AAAUUUUAAAAAAAAAU | 17 |
| AAAAAUUUUUUAAAAAU | 17 |
| UAAAUUUAAAAAAAC | 15 |
| AAUUAUUAAAAAAU | 14 |
| AUUCUUUAAAAAC | 13 |
| AAAUAAAAAAAC | 12 |
| AAAAAUAAAAAU | 12 |
| AAAAUUAAAU | 10 |
| UAUAAAU | 7 |
| UUAAAAU | 7 |
| AAAAAC | 6 |
| UAAAU | 5 |
| AAAAU | 5 |
| AAAC | 4 |
| AAU | 3 |
| AAU | 3 |
| ACU | 3 |
| average: | 11.95652174 |
| standard deviation: | 6.724975572 |

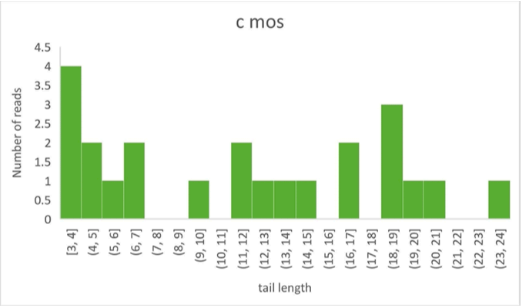

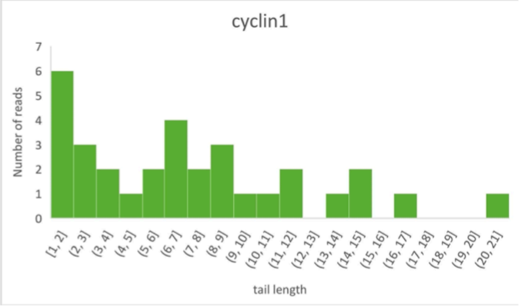

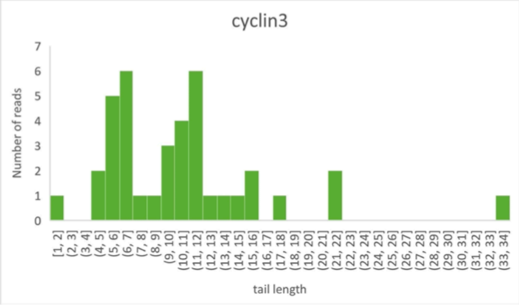

**Supplementary Figure S8.** Non-templated sequence in the 3'-end of *N. vectensis* egg mRNA reads

(NCBI BioProject: PRJNA956458 obtained using Oxford Nanopore Technology Direct RNA sequencing methodology. Genic sequence for *cyclin1*, *cyclin3*, and *c-mos* was recognized by sequence alignment and the remainder of the 3'end of individual reads (after removal of adapter oligo sequence, when present) is shown for each gene.  Graphs summarizing the observed length of tails using this method is show for each of the three genes.

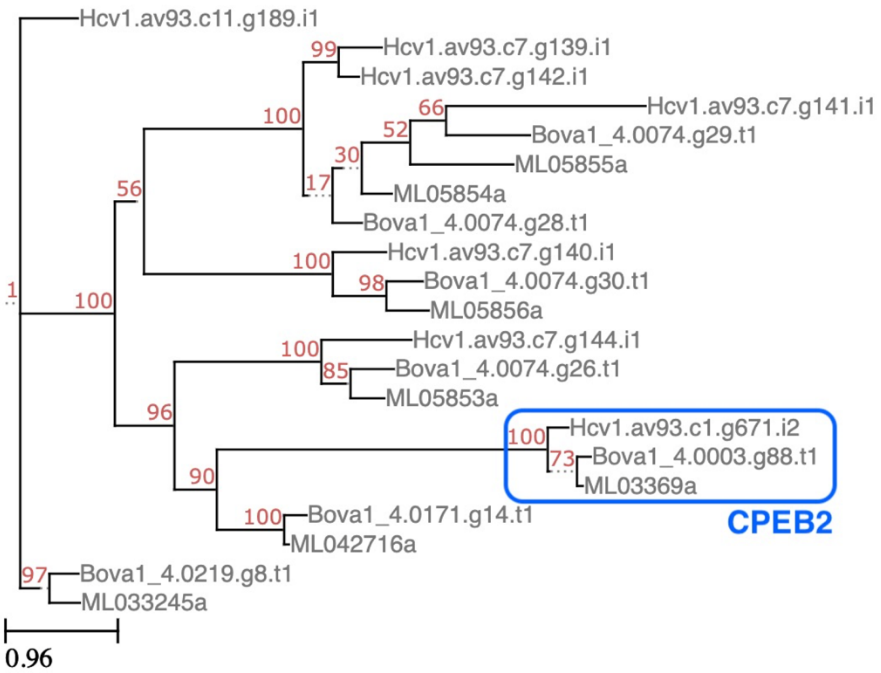

**Supplementary Figure S9**. Phylogram showing position of CPEBs within Ctenophora. CPEBs from *Hormiphora californensis* and *Beroe ovata* were identified via reciprocal BLAST between human CPEB protein sequences (CPEB1, NP_001275748.1; and CPEB4, AAH36899.1) and the protein models of these two ctenophore species and aligned with *Mnemiopsis leidyi* CPEBs using MAFFT (default parameters). The position of CPEBs was calculated according to maximum-likelihood analysis using IQ-TREE (default parameters with automatic model finding and 1000 bootstraps) and the phylogram generated using ETE toolkit (http://etetoolkit.org/; [175]). Scale bar represents substitutions per amino acid position.

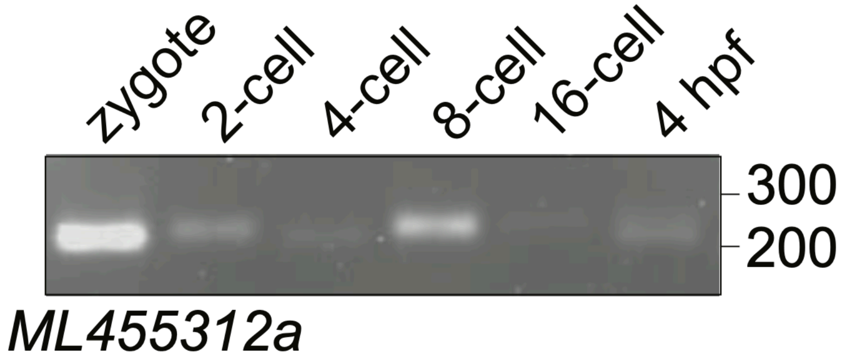

**Supplementary Figure S10**. Detailed analysis of changes in poly(A) tail length using the method described in Figure 2K for *M. leidyi cyclin* homolog *ML455312a* during each of the first four cleavages and at 4-hpf. Position of DNA size markers is shown on the right.

### Supplementary Tables

**Supplementary Table S1**. CPEBs orthologs found in genomes of 44 animal species, including representatives from 14 animal phyla, but not in genomes from species outside the Metazoa (modified from Paps and Holland [80].

| | |
|---|---|
| **Deuterostomia** | |
| *Saccoglossus kowalevskii* | Skow_10916m Skow_22287m |
| *Strongylocentrotus purpuratus* | Spur_015450_Cpeb1 Spur_001293_Cpeb2 |
| *Branchiostoma floridae* | Bflo_260831047 |
| *Ciona intestinalis* | Cint_003938 Cint_024148 Cint_003274 |
| *Ciona savignyi* | Csav_004444 Csav_013417 Csav_004446 Csav_004445 Csav_013416 Csav_013415 |
| *Botryllus schlosseri* | Bsch_g10689 Bsch_g8517 Bsch_g25882 Bsch_g45408 Bsch_g63388 |
| *Oikopleura dioica* | Odio_015065001 |
| *Danio rerio* | Drer_008454_cytoplasmic_polyadenylation_element_binding_protein_1b_Source_ZFIN_Acc_ZDB_GENE_990927_1_cpeb1b<br>Drer_053117_Uncharacterized_protein_Source_UniProtKB_TrEMBL_Acc_E7FA11_cpeb1a<br>Drer_045932_cytoplasmic_polyadenylation_element_binding_protein_1a_Source_ZFIN_Acc_ZDB_GENE_080723_42_cpeb1a<br>Drer_056691_cytoplasmic_polyadenylation_element_binding_protein_4_Source_ZFIN_Acc_ZDB_GENE_040426_1557_cpeb4<br>Drer_060103_cytoplasmic_polyadenylation_element_binding_protein_3_Source_ZFIN_Acc_ZDB_GENE_090312_68_cpeb3<br>Drer_052604_cytoplasmic_polyadenylation_element_binding_protein_2_Source_HGNC_Symbol_Acc_21745_CPEB2<br>Drer_074474_cytoplasmic_polyadenylation_element_binding_protein_4_Source_HGNC_Symbol_Acc_21747_CPEB4_3_of_3<br>Drer_060495_cytoplasmic_polyadenylation_element_binding_protein_4_Source_HGNC_Symbol_Acc_21747_CPEB4_2_of_3 |
| *Xenopus (Silurana) tropicalis* | Xtro_023343 Xtro_059795 Xtro_026614 Xtro_043840 Xtro_027833 |
| *Gallus gallus* | Ggal_003366 Ggal_011187 Ggal_023384 Ggal_022748 |
| *Anolis carolinensis* | Acar_00000309 Acar_00014820 Acar_00020487 Acar_00006260 Acar_00011854 |
| *Homo sapiens* | Hsap_Ensmbl0260836_RP11_152F13_10<br>Hsap_Ensmbl0214575_cytoplasmic_polyadenylation_element_binding_protein_1_Source_HGNC_Symbol_Acc_21744_CPEB1<br>Hsap_Ensmbl0107864_cytoplasmic_polyadenylation_element_binding_protein_3_Source_HGNC_Symbol_Acc_21746_CPEB3<br>Hsap_Ensmbl0113742_cytoplasmic_polyadenylation_element_binding_protein_4_Source_HGNC_Symbol_Acc_21747_CPEB4<br>Hsap_Ensmbl0137449_cytoplasmic_polyadenylation_element_binding_protein_2_Source_HGNC_Symbol_Acc_21745_CPEB2 |
| **Ecdysozoa** | |
| *Trichinella spiralis* | Tspi_316973214 Tspi_316969994 Tspi_316978840 Tspi_316956776 |
| *Romanomermis culicivorax* | Rcul_t24732 Rcul_t40503 Rcul_t00011 Rcul_t00012 Rcul_t01824 |
| *Caenorhabditis elegans* | Cele_000772_cpb_3_Protein_CPB_3_cpb_3_mRNA_complete_cds_Source_RefSeq_mRNA_Acc_NM_059279<br>Cele_000770_cpb_1_Protein_CPB_1_cpb_1_mRNA_complete_cds_Source_RefSeq_mRNA_Acc_NM_066650<br>Cele_001481_fog_1_Protein_FOG_1_isoform_b_Source_RefSeq_mRNA_Acc_NM_001026621<br>Cele_000771_cpb_2_Protein_CPB_2_cpb_2_mRNA_complete_cds_Source_RefSeq_mRNA_Acc_NM_062835 |
| *Brugia malayi* | Bmal_170571749 Bmal_170579891 Bmal_170596242 Bmal_170582725 |
| *Strigamia maritima* | Smar_004137 Smar_004113 |
| *Mesobuthus martensii* (scorpion) | Mmar_MMa39152 Mmar_MMa28869 Mmar_MMa28867 Mmar_MMa45969 |
| *Stegodyphus mimosarum* (spider) | Smim_72406_cytoplasmic_polyadenylation_element_binding_protein_putative_Ixodes_scapularis_gi_241829409_ref_XP_0024 14761_6e_175_501<br>Smim_56756_PREDICTED_probable_RNA_binding_protein_orb2_like_isoform_Acyrthosiphon_pisum_gi_328721760_ref_XP_ 003247398_0_0_575<br>Smim_27690_cytoplasmic_polyadenylation_element_binding_protein_putative_Ixodes_scapularis_gi_241560350_ref_XP_0024 00996_2e_85_260<br>Smim_27683_A_Chain_A_Solution_Structure_Of_Rna_Binding_Domain_In_Cytoplasmic_Polyadenylation_Element_Binding_Pr otein_3_gi_159164253_pdb_2DNL_A_5e_40_136 |
| *Ixodes scapularis* (tick) | Isca_ISCW023425-PA Isca_ISCW010113-PA |
| *Daphnia pulex* | Dpul_326182 Dpul_308504 |
| *Zootermopsis nevadensis* | Znev_12790 Znev_18107 |
| *Tribolium castaneum* (beetle) | Tcas_270001277 Tcas_270015159 |
| *Drosophila melanogaster* | Dmel_0004882_orb_oo18_RNA_binding_protein_Source_FlyBase_0004882 Dmel_0264307_orb2 |
| **Lophotrochozoa** | |
| *Adineta vaga* | Avag_50023001 Avag_12615001 Avag_47368001 Avag_01285001 Avag_12616001 Avag_49767001 Avag_04121001<br>Avag_07376001 Avag_00865001 Avag_15043001 Avag_65428001 Avag_61392001 Avag_18987001 Avag_04122001<br>Avag_11453001 Avag_59612001 Avag_19009001 Avag_27103001 Avag_50024001 Avag_01286001 Avag_49768001 |
| *Gyrodactylus salaris* | Gsal_4058_cytoplasmic_polyadenylation_element_binding_protein_1<br>Gsal_1072_cytoplasmic_polyadenylation_element_binding_protein_2 |
| *Schistosoma japonicum* | Sjap_0065230 Sjap_0032030 Sjap_0029950 |

| | |
|---|---|
| *Schistosoma mansoni* | Sman_070360_Putative_cytoplasmic_polyadenylation_element_binding_protein_Cpeb_Source_UniProtKB_TrEMBL_Acc_G4VT97 Sman_012950_Putative_cytoplasmic_polyadenylation_element_binding_protein_Cpeb_Source_UniProtKB_TrEMBL_Acc_G4VEX4 Sman_136410_Putative_cytoplasmic_polyadenylation_element_binding_protein_Cpeb_Source_UniProtKB_TrEMBL_Acc_G4VEX5 Sman_137460_Cytoplasmic_polyadenylation_element_binding_protein_Cpeb_putative_Source_UniProtKB_TrEMBL_Acc_G4LW19 |
| *Hymenolepis microstoma* | Hmic_0079100 |
| *Echinococcus granulosus* | Egra_08380 Egra_00993 |
| *Echinococcus multilocularis* | Emul_08189 Emul_03389 |
| *Crassostrea gigas* | Cgig_14557_Cytoplasmic_polyadenylation_element_binding_protein_4_Source_UniProtKB_TrEMBL_Acc_K1Q726 Cgig_19114_Cytoplasmic_polyadenylation_element_binding_protein_1_B_Source_UniProtKB_TrEMBL_Acc_K1QYV5 |
| *Pinctada fucata* | Pfuc_154180_1_28450_t1 Pfuc_1867_1_37153_t1 Pfuc_36185_1_26743_t1 |
| *Lottia gigantea* | Lgig_213887 Lgig_115324 |
| *Capitella teleta* | Ctel_118515 Ctel_103382 |
| *Helobdella robusta* | Hrob_111676 Hrob_79994 Hrob_69117 Hrob_66900 |
| **Cnidaria** | |
| *Nematostella vectensis* | Nvec_39034_ Nvec_34062_ Nvec_57237_RRM_1 |
| *Acropora digitifera* | Adig_13400 Adig_10936 |
| *Hydra magnipapillata* | Hmag_224246 Hmag_210731 Hmag_202650 |
| **Placozoa** | |
| *Trichoplax adhaerens* | Tadh_30543 Tadh_58533 |
| **Porifera** | |
| *Amphimedon queenslandica* | Aque_228234 Aque_215503 |
| *Oscarella carmela* (incomplete?) | OcarG_3927 OcarG_1845 |
| **Ctenophora** | |
| *Mnemiopsys leidyi* | Mley_ML033245a Mley_ML05854a Mley_ML05855a Mley_ML05856a Mley_ML03369a Mley_ML042716a |
| *Pleurobrachia bachei* | Pbac_3462211 Pbac_3463340 Pbac_3460394 Pbac_3479762 Pbac_3475771 |
| **Non-Metazoan Eukaryotes** | |
| *Chlamydomonas reinhardtii* | |
| *Physcomitrella patens* | |
| *Selaginella moellendorffii* | |
| *Amborella trichopoda* | |
| *Arabidopsis thaliana* | |
| *Emiliana huxleyii* | |
| *Bigelowiella natans* | |
| *Reticulomyxa filosa* | |
| *Thecamonas trahens* ATCC 50062 | |
| *Fonticula alba* | |
| *Spizellomyces punctatus* | |
| *Allomyces macrogynus* | |
| *Saccharomyces cerevisiae* | |
| *Sphaeroforma arctica* | |
| *Creolimax fragrantissima* | |
| *Capsaspora owczarzaki* | |
| *Monosiga brevicollis* | |
| *Salpingoeca rosetta* | |

**Supplementary Table S2**. Percent nucleotide composition of 5' halves, 3' halves, and full poly(A) tails obtained from Amplicon-EZ reads of *N. vectensis c-mos, cyclin 1*, and *cyclin 2* mRNAs. Average from all three mRNAs including both developmental stages tested for each is shown at the bottom.

| ***c-mos*** **(oocyte)** | **5' half** | **3' half** | **total** |
|---|---|---|---|
| **A** | 75 | 92 | **83.5** |
| **U** | 12 | 4 | **8** |
| **C** | 5 | 1 | **3** |
| **G** | 9 | 3 | **6** |

| **c-mos (egg)** | | | |
|---|---|---|---|
| **A** | 69 | 92 | **80.5** |
| **U** | 24 | 6 | **15** |
| **C** | 3 | 1 | **2** |
| **G** | 4 | 1 | **2.5** |

| ***cyclin 1*** **(oocyte)** | | | |
|---|---|---|---|
| **A** | 42 | 80 | **61** |
| **U** | 39 | 14 | **26.5** |
| **C** | 6 | 1 | **3.5** |
| **G** | 12 | 5 | **8.5** |

| **cyclin 1 (egg)** | | | |
|---|---|---|---|
| **A** | 63 | 94 | **78.5** |
| **U** | 30 | 4 | **17** |
| **C** | 3 | 0 | **1.5** |
| **G** | 4 | 1 | **2.5** |

| ***cyclin 3*** **(egg)** | | | |
|---|---|---|---|
| **A** | 55 | 86 | **70.5** |
| **U** | 27 | 7 | **17** |
| **C** | 7 | 3 | **5** |
| **G** | 10 | 4 | **7** |

| **cyclin 3 (2 HPF)** | | | |
|---|---|---|---|
| **A** | 71 | 92 | **81.5** |
| **U** | 22 | 6 | **14** |
| **C** | 5 | 2 | **3.5** |
| **G** | 3 | 1 | **2** |

| **Average (All)** | **5' half** | **3' half** | **total** |
|---|---|---|---|
| **A** | 62.50 | 89.33 | **75.92** |
| **U** | 25.67 | 6.83 | **16.25** |
| **C** | 4.83 | 1.33 | **3.08** |
| **G** | 7.00 | 2.50 | **4.75** |

**Supplementary Table S3.** Identification of CPEB homologs in three different ctenophores. CPEB1 and CPEB2 orthologs identified in reference sequences (left column) of *Mnemiopsis leidyi* (top), *Hormiphora californensis* (center), and *Beroe ovata* (bottom). Transcripts or gene models concluded to correspond to individual genes are grouped in individual rows. Top match from reciprocal BLASTP analyses with human proteins and corresponding E-values (center columns) are included. Glutamine (Q)-rich regions identified as included in at least three of four consecutive positions are indicated (right column).

| | Transcript ID | Top Hs Match (RefSeq proteins) | BLASTP E-val (vs. Hs Match) | Potential Prion Motifs (minimum 3 Q in 4 AA stretch) |
|---|---|---|---|---|
| **M. leidyi CPEBs (TBLASTN vs. NHGRI Gene models 2.2)** | Ml-CPEB1a (ML042716) | CPEB1 | 1E-129 | QQQQQPQ; QVPQQSQPDQ |
| | Ml-CPEB1b (ML05853a) | CPEB1 | 1E-71 | - |
| | Ml-CPEB1c (ML05854a) | CPEB1 | 1E-60 | - |
| | Ml-CPEB1d (ML05855a) | CPEB1 | 4E-54 | - |
| | Ml-CPEB1e (ML05856a) | CPEB1 | 2E-61 | - |
| | *Ml-CPEB1f* (ML033245a) | CPEB1 | 6E-78 | - |
| | *Ml-CPEB2* (ML03369a) | CPEB4 | 6E-142 | |
| **TBLASTN vs. Hcv1av93_transcripts .fasta** | Hcv1.av93.c11.g189.i1<br>Hcv1.av93.c11.g188.i1 | CPEB1 | 7E-72<br>8E-68 | - |
| | Hcv1.av93.c7.g144.i1 | CPEB1 | 1E-63 | - |
| | Hcv1.av93.c7.g142.i1 | CPEB1 | 1E-60 | QQQMTPSLQLQ; QKQQQYHPQQHSQQHSQQQQ |
| | Hcv1.av93.c7.g139.i1 | CPEB1 | 3E-54 | - |
| | Hcv1.av93.c7.g141.i1 | CPEB1 | 2E-40 | - |
| | Hcv1.av93.c7.g140.i1 | CPEB1 | 5E-39 | QQQQQPQQ; QQSQYQQQ; QEQQ; QNKQQPQAAPTQQQQ |
| | Hcv1.av93.c1.g671.i2<br>Hcv1.av93.c1.g671.i1<br>Hcv1.av93.c1.g671.i3 | CPEB4 | 8E-140<br>7E-135<br>4E-131 | QQQHQQQ; N-rich |
| **TBLASTN vs. Bova1.4 cds** | Bova1_4.0171.g14.t1 | CPEB1 | 2E-107 | QQQ; QPQPQQHQ; QQQPPPQQQPQQPPQ |
| | Bova1_4.0219.g8.t1 | CPEB1 | 5E-74 | - |
| | Bova1_4.0074.g29.t1 | CPEB1 | 4E-77 | - |
| | Bova1_4.0074.g26.t1 | CPEB1 | 3E-64 | - |
| | Bova1_4.0074.g28.t1 | CPEB1 | 5E-59 | - |
| | Bova1_4.0074.g29.t1 | CPEB1 | 1E-47 | - |
| | Bova1_4.0003.g88.t3<br>Bova1_4.0003.g88.t4<br>Bova1_4.0003.g88.t2<br>Bova1_4.0003.g88.t1 | CPEB3 | 9E-137<br>2E-135<br>4E-134<br>5E-134 | - |

**Supplementary Files (available at https://github.com/lrouhana/cpeb_evolution)**

**Supplementary File S1.** Protein sequences with homology to human CPEB1 that were identified in BLAST searches against reference sequences from animal and non-animal species. Redundant sequences were removed and only longer isoforms were retained.

**Supplementary File S2.** Amplicon-EZ reads of 3'ends of *Nv_c-mos*, *Nv_cyclin 1*, and *Nv_cyclin 3* mRNAs from stages analyzed in Figure 2 M-O and Supplementary Figure S6.

**Supplementary File S3.** *N. vectensis* egg transcriptome sequences generated using Oxford Nanopore Direct RNA-sequencing (available for download under NCBI BioProject: PRJNA956458).
